# Genomic insights into variation in thermotolerance between hybridizing swordtail fishes

**DOI:** 10.1101/2021.10.22.465484

**Authors:** Cheyenne Payne, Stephen Bovio, Daniel Powell, Theresa Gunn, Shreya Banerjee, Victoria Grant, Gil Rosenthal, Molly Schumer

**Affiliations:** Department of Biology, Stanford University, Stanford, CA, USA; Centro de Investigaciones Científicas de las Huastecas “Aguazarca”, A.C., Calnali, Hidalgo, México; Department of Biology, Texas A&M University; Hanna H. Gray Fellow, Howard Hughes Medical Institute, Stanford, CA, USA

**Keywords:** thermotolerance, hybridization, swordtail fishes, misexpression, molecular ecology

## Abstract

Understanding how organisms adapt to changing environments is a core focus of research in evolutionary biology. One common mechanism is adaptive introgression, which has received increasing attention as a potential route to rapid adaptation in populations struggling in the face of ecological change, particularly global climate change. However, hybridization can also result in deleterious genetic interactions that may limit the benefits of adaptive introgression. Here, we used a combination of genome-wide quantitative trait locus mapping and differential gene expression analyses between the swordtail fish species *Xiphophorus malinche* and *X. birchmanni* to study the consequences of hybridization on thermotolerance. While these two species are adapted to different thermal environments, we document a complicated architecture of thermotolerance in hybrids. We identify a region of the genome that contributes to reduced thermotolerance in individuals heterozygous for *X. malinche* and *X. birchmanni* ancestry, as well as widespread misexpression in hybrids of genes that respond to thermal stress in the parental species, particularly in the circadian clock pathway. We also show that a previously mapped hybrid incompatibility between *X. malinche* and *X. birchmanni* contributes to reduced thermotolerance in hybrids. Together, our results highlight the challenges of understanding the impact of hybridization on complex ecological traits and its potential impact on adaptive introgression.

## Introduction

Hybridization, or interbreeding between species, is much more common than previously thought and can have diverse genetic and evolutionary consequences [1]. For example, a large body of work has shown that hybridization can facilitate the movement of adaptive alleles between species, promoting ecological adaptation to novel or changing environments [2–16]. In hybridizing species, gene flow may serve as a mechanism of rapid adaptation [7,8,16,17], since adaptive introgression can occur on a shorter timescale than that required for new adaptive mutations to arise within a species [18,19].

While hybridization has played a role in adaptation on evolutionary timescales [19–21], hybridization is thought to occur more frequently under environmental disturbance and stress [12,13,22–24]. As environmental conditions shift due to climate change, understanding the genetic mechanisms that can facilitate rapid adaptation will be critical in predicting whether vulnerable populations will adapt or collapse [18,25]. This in turn requires characterizing the genetic architecture of ecologically relevant traits that distinguish hybridizing species [26–29].

Although there are many examples of adaptive introgression between species [7–9,18], deleterious effects of hybridization are also well-documented and have been studied for decades [30,31]. Hybridization frequently uncovers negative interactions between mutations that have arisen independently in the genomes of the two parental species. These interactions can result in reduced hybrid viability or fertility [32–34], and their costs may outweigh the potential benefits of hybridization as a source of adaptive alleles [1]. Such interactions are commonly known as Bateson-Dobzhansky-Muller incompatibilities (BDMIs; [35]). While BDMIs were originally envisioned to result from incompatible interactions between proteins encoded by two or more genes, recent work has highlighted the diversity of mechanisms through which BDMIs may arise [36–42]. Recently attention has been paid to regulatory BDMIs, which arise from the coevolution of *cis* and *trans* regulatory elements within species that become mismatched and cause misexpression of target genes in hybrids [1,36,43,44] (here, misexpression is defined as expression of genes in hybrids that is much higher or lower than that observed in either parent species).

BDMIs can impact a range of traits, including those relevant for an individual’s survival in their environment [45,46]. In fact, both theory and empirical results suggest that BDMIs may frequently arise from divergent adaptation to the environment [47–49]. In addition to the expectation that hybrids may have reduced ecological fitness due to phenotypic intermediacy or dominance of particular parental traits [45,46,50–54], BDMIs can arise at loci underlying ecological traits. Despite their predicted importance, few ecological BDMIs have been identified to date [46] (see [55] and [46] for examples from *Arabidopsis* and sticklebacks), making it difficult to study the tradeoffs between adaptive introgression of ecological traits and selection on ecological BDMIs in hybrids.

An ecological trait that can be used to address this gap and that is of particular interest for predicting how populations may adapt to global climate change is thermal tolerance [56–58]. Though thermal tolerance can be defined in many ways, as global temperatures warm, a relevant thermal tolerance trait is an organism’s upper thermal limit (hereafter referred to as “thermotolerance”) [59,60]. Little work to date has explored whether loci that control variation in thermotolerance introgress between hybridizing animal populations or how effective such introgression is as a mechanism of thermal adaptation (though suggestive results have been reported in some species; box turtles [61]; wasps [12]; copepods [62,63]). The deficit in empirical work in this area is likely due to the paired difficulties of mapping the genetic basis of complex traits like thermotolerance and studying them in natural hybrid populations.

Here, we take advantage of a system where natural hybridization is ongoing between species that vary in thermotolerance. Two sister species of swordtail fishes, *Xiphophorus birchmanni* and *X. malinche*, are endemic to rivers in eastern México [64], and their distributions are determined in part by their thermal habitats [65]. *X. malinche* lives in cooler (7-25°C) streams at high elevations, while the more heat tolerant *X. birchmanni* lives downstream in the warmer lowlands (15-35°C; [64]). These species are sympatric in regions where their temperature ranges overlap. Recently, pollution has interfered with species-specific olfactory communication, causing breakdown of mating barriers [66,67]. As a result, natural hybrid zones have formed, with clinal ancestry patterns that mirror thermal gradients, where there is low *X. birchmanni* ancestry and low thermotolerance in the highlands and high *X. birchmanni* ancestry and high thermotolerance in the lowlands [64,65]. Though phenotypic plasticity contributes to this differential tolerance (as shown in [65]), we show here that variation in innate thermotolerance between species is in part genetic. Leveraging this finding, we combine thermotolerance assays with classic quantitative trait locus (QTL) mapping, gene expression analysis, and analysis of ancestry in natural hybrid populations to explore the evolution of thermotolerance in this system. Unexpectedly, we find that individuals that are heterozygous for ancestry in one genomic region have reduced critical thermal maxima, and F_1_ hybrids have widespread misexpression of core regulatory genes of the circadian clock, which appear to be associated with proper thermal regulation. Additionally, we uncover a relationship between reduced thermotolerance and a previously mapped hybrid incompatibility.

## Methods

### Comparison of CT_max_ between X. malinche, X. birchmanni, F_1_s, and F_2_s and measurement of CT_max_ for QTL mapping

One ecologically-relevant measure of upper thermotolerance in ectotherms is the critical thermal maximum, or CT_max_ [59]. Specifically, the CT_max_ of a fish is the highest temperature it can withstand before it experiences loss of equilibrium and is unable to maintain its balance [59,60]. We tested CT_max_ for *Xiphophorus malinche, X. birchmanni*, and F_1_ and F_2_ hybrids between the two species reared in a common garden environment.

We simultaneously reared *X. malinche* fry born to wild-caught mothers from the Chicayotla locality on the Río Xontla (1003 meters elevation; 20°55’27.24”N 98°34’34.50W), *X. birchmanni* fry from wild-caught mothers from the Coacuilco locality on the Río Coacuilco (320 meters elevation; 21°5’50.85 N, 98°35’19.46 W), and F_1_ and F_2_ fry generated from these parent populations (Fig. 1A). Specifically, F_1_ fry were generated by crossing *X. malinche* (Chicayotla) females to *X. birchmanni* (Coacuilco) males, and F_2_ fry were generated by intercrossing previously produced F_1_s (Fig. 1B). We note that due to the crossing design, all artificial hybrids in this study harbor *X. malinche* mitochondria; crosses in the reverse direction are largely unsuccessful. All fish were crossed and raised in 2,000 L semi-natural mesocosms at the CICHAZ field station in Calnali, Hidalgo, México. Individuals for all four groups were born between 16 May and 24 May 2016, at which time offspring from each group were randomly assigned to one of three replicate 2,000 L semi-natural mesocosms for a total of 12 tanks (three per class, n = 34 per tank).

**Figure 1.**
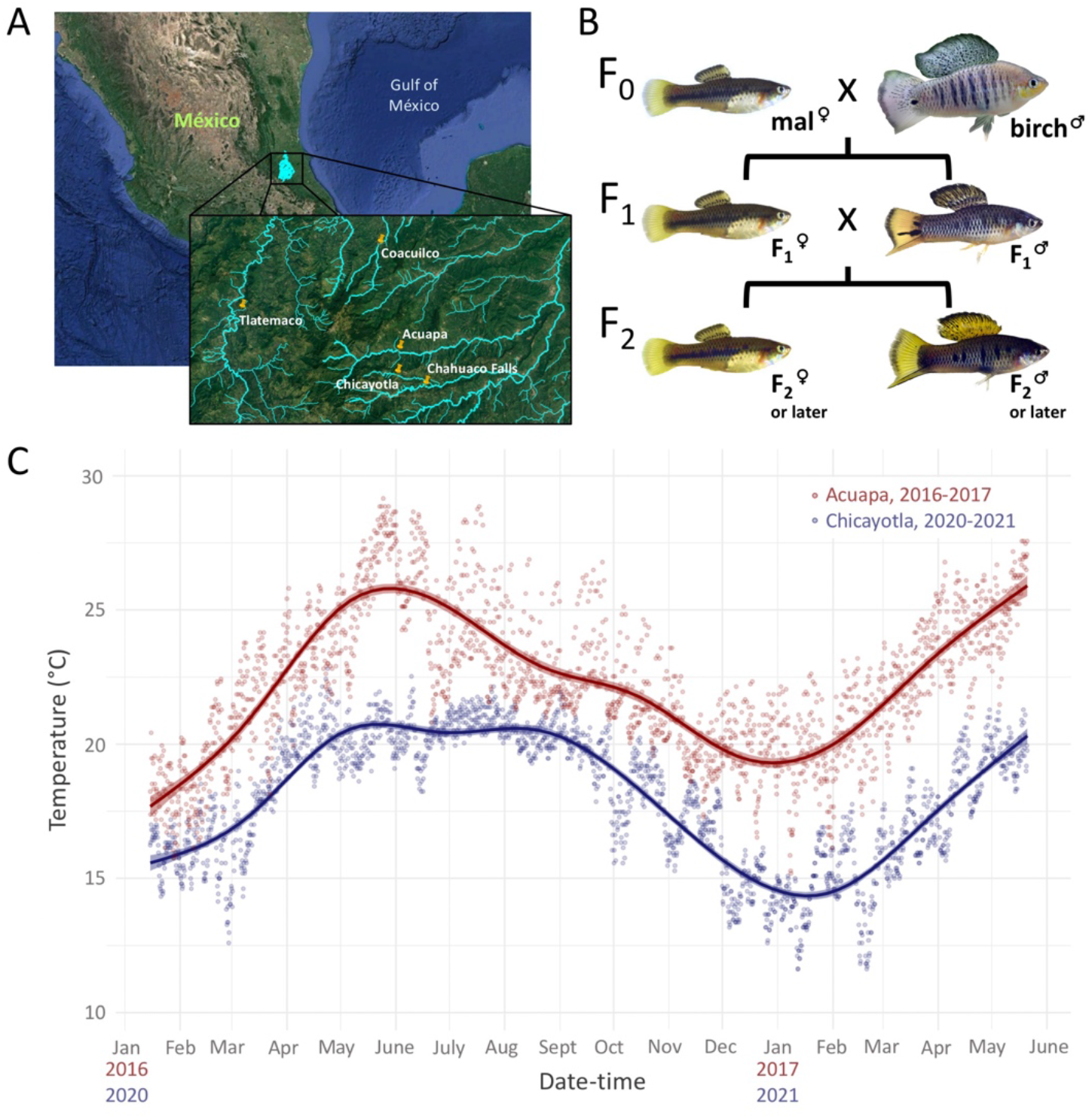
**A**. Map adapted from Google Earth showing the five natural populations from which fish were collected for data used in this study. Pure *X. malinche* mothers and *X. birchmanni* fathers used in crosses and for RNAseq experiments were originally collected from the Chicayotla and Coacuilco populations, respectively. Natural hybrids were collected from Chahuaco Falls to evaluate links between hybrid melanoma and CT_max_, and natural hybrids for analysis of population-level ancestry were collected from the low elevation Acuapa and Tlatemaco hybrid populations. **B**. The cross design used to generate individuals for both the mapping and RNAseq datasets. Wild *X. malinche* mothers from Chicayotla and *X. birchmanni* fathers from Coacuilco were crossed create an F_1_ population. A subset of F_1_s were crossed to generate an artificial hybrid mapping population that was raised in common garden conditions. Other F_1_ individuals were raised in the lab and used for the RNAseq thermal stress experiment. Abbreviations: mal – *X. malinche*, birch – *X. birchmanni*. **C**. Temperature data collected by HOBO loggers deployed at Acuapa from 2016-2017 and Chicayotla from 2020-2021. Acuapa is a hybrid population that is found at a similar elevation to pure *X. birchmanni* sites (∼400 meters versus ∼250-300 meters; [64]), and Chicayotla is a site where pure *X. malinche* individuals are found (∼1000 meters). Data points were collected four times per day by the loggers. Points and trend lines are shown in red for Acuapa and blue for Chicayotla.

To measure variation in CT_max_ between *X. malinche, X. birchmanni*, F_1_s, and F_2_s, CT_max_ trials were performed in February 2018 using methods similar to Culumber et al [65]. Trials followed procedures approved in Texas A&M IACUC protocol #117419. Briefly, the test fish (eight per trial, mix of males and females from one group), an air bubbler, a standard glass thermometer, and a HOBO temperature logger (Onset) were placed in an enamel container holding 4 L of water at ambient temperature (16.1 ± 0.2°C). The enamel container was nested in a larger container of water which was suspended above a hot plate. Water was heated at a standardized ramp-up rate of 0.3 °C/min until the fish lost equilibrium (following Becker & Genoway [60]). The time and temperature of initial loss of equilibrium (i.e. the first time balance is lost) for each fish was recorded, and the fish was immediately placed in an ambient temperature recovery tank. Because the data departed from the assumption of normality, we used a Mann-Whitney Wilcoxon test to evaluate the effect of genotype on CT_max_ (Table S1).

We repeated these procedures for a larger mapping population of 152 *X. malinche-X. birchmanni* artificial hybrids. Due to the difficulty of raising sufficient numbers of individuals in common garden conditions, our mapping population included individuals ranging from F_2_-F_4_ generations, initially generated from F_1_ intercrosses. For each individual, we collected a fin clip from each fish at the end of the CT_max_ trial to perform QTL mapping. In addition to CT_max_ time and temperature, we recorded metadata for each fish to account for potential covariates in mapping. We found that one of the strongest covariates with CT_max_ was rearing tank number (which also corresponded to trial number). Therefore, we combined these covariates into a single variable that we refer to as tank throughout the manuscript (listed as ‘site.tank’ in data files).

### DNA extraction and library preparation

Fin clips were added to 96 well plates and DNA was extracted using the Agencourt DNAdvance bead-based kit. The protocol followed that specified by the manufacturer except that we used half-reactions. We quantified extracted DNA using a TECAN microplate reader. After diluting DNA to 2.5 ng/ul, we prepared tagmentation-based libraries for low-coverage whole genome sequencing. DNA was enzymatically sheared using the Illumina Tagment DNA TDE1 Enzyme and Buffer Kits by incubating DNA, buffer and enzyme at 55°C for 5 minutes. Fragmented DNA was amplified in a dual-indexed PCR reaction for 12 cycles and PCR-products were pooled and bead purified with 18% SPRI magnetic beads. Purified libraries were quantified using a Qubit fluorometer and library size distribution was evaluated using an Agilent 4200 Tapestation.

### Artificial hybrid QTL mapping sample sequencing and genotyping

Low-coverage whole genome sequence data was collected from these libraries on an Illumina 4000 machine using 150 bp paired-end reads (∼0.1-0.3X per basepair coverage). Using the program *ancestryinfer* [68], reads were mapped to both the *X. birchmanni* and *X. malinche* genomes with BWA-MEM [69], and those that showed evidence of mapping bias or that did not map uniquely were discarded. Reads matching each parental allele at ancestry-informative sites were counted from samtools mpileup files [70], and informative sites were thinned to one per read to minimize errors due to mismapping. This data was input into AncestryHMM [71], a hidden Markov model (HMM) based local ancestry inference program that relies on read counts at ancestry informative sites and transition probabilities to infer posterior probabilities for ancestry states (in our case: homozygous *X. birchmanni*, heterozygous, or homozygous *X. malinche*). Past work has shown that this low-coverage whole genome sequencing approach has excellent accuracy for early generation *X. malinche x X. birchmanni* hybrids [68,72]. This analysis yielded posterior probabilities for each ancestry state at ∼700,000 ancestry-informative sites across the genome (approximately one per kb).

Because it was convenient for downstream analyses, we converted posterior probabilities at each ancestry informative site to hard genotype calls. For each sample, markers with greater than 0.9 posterior probability for any ancestry state were assigned to that state; markers with less than 0.9 posterior probability for any ancestry state were converted to NAs. Homozygous *X. birchmanni*, heterozygous, and homozygous *X. malinche* ancestry calls were assigned genotypes of 0, 1, and 2 respectively.

### CT_max_ QTL mapping analysis

To identify regions of the genome that are associated with variation in thermotolerance, we used a QTL mapping approach. We performed QTL mapping with R/qtl [73] to scan for an association between genotypes at ancestry-informative markers across the genome and the CT_max_ phenotype. For computational efficiency, markers were thinned to retain at most one marker per 20 kb. Ancestry linkage disequilibrium decays over several megabases in early generation hybrids [74]; thus, we do not expect to sacrifice any power to map QTL by performing this thinning. The thinning step resulted in 30,244 ancestry informative markers retained throughout the genome, or ∼45 per Mb.

Data were converted to the R/qtl input format using custom scripts (https://github.com/Schumerlab/thermotolerance). Input files for analysis with R/qtl included CT_max_, covariate data (e.g. tank), and genotype data for all 152 individuals. Markers with fewer than 80% of individuals genotyped (i.e. less than 120 out of 152) were filtered. Several individuals had high levels of missing data (>25% of markers with an NA ancestry state) and these individuals were removed. After filtering, 144 individuals and 29,652 markers were retained, with ∼95% of individuals genotyped at any given marker. Next markers were evaluated for segregation distortion at a Bonferroni corrected p-value < 0.05 using R/qtl’s internal commands, and 610 markers on chromosome 13 were dropped, resulting in 29,042 markers for the QTL scan. Recombination frequency and genotype probabilities were calculated using the `est.rf` and `calc.genoprob` functions, respectively.

To select an appropriate model for mapping in R/qtl, we used the R step function to calculate AIC for models incorporating a suite of possible covariates, including the tank variable (tank), hybrid index (the proportion of the genome derived from the *X. malinche* parent), genome-wide ancestry heterozygosity, and sex (e.g. CT_max_ ∼ hybrid_index + heterozygosity + tank + sex). We retained all tank variables with a significant effect on CT_max_ (17 total) and used a method called ‘one-hot encoding’ to recode the tank variable so that the tank variable would be treated categorically by R/qtl; other covariates were not retained in the step function analysis. Even though hybrid index was not retained, we included it in our final mapping model since past work has suggested that failing to include ancestry as a covariate can result in artifacts in QTL analysis [72].

A genome-wide scan with a single-QTL model was performed with the `scanone` function, using the Haley-Knott regression method [75] and the tank and hybrid index covariates, as described above. The 5% and 10% false discovery rate thresholds were estimated with 1,000 permutations (LOD thresholds of 4.72 and 4.33 respectively), where CT_max_ phenotypes were shuffled onto genotypes and a QTL scan conducted 1,000 times to create a null distribution of associations expected by chance. To search for potential interacting QTL, we performed a second scan using the same method, but added genotypes at the chromosome 22 QTL peak as an interaction term in the model (significant thresholds at the 5% and 10% FDR level for the interaction analysis were 9.63 and 8.96, respectively).

### Estimating the effect size of detected QTL

We identified one QTL on chromosome 22 and one putative interacting QTL on chromosome 15 that were significantly associated with variation in CT_max_ after controlling for other covariates at the 10% false discovery threshold (see Results). We used two methods to obtain estimates for the effect sizes of these QTL (i.e. the percentage of the variation in CT_max_ explained by each QTL and their interaction). First, we used the drop-one-term analysis from fitting a multiple QTL model with the R/qtl function `fitqtl`, to estimate the effect sizes of the chromosome 22 and 15 QTL on CT_max_, as well as to estimate the effect size of their interaction. Because effect size estimates are often inflated in QTL studies with low statistical power [76], we also performed simulations to explore the range of possible effect sizes consistent with our empirical results for the main effect QTL on chromosome 22. Methods and results for those simulations are reported in Supporting Information 1.

### Multiple tissue thermal stress RNAseq experiment, library preparation, and sequencing

To compare expression of genes across the genome that respond to high temperature in the two parental species and their hybrids, we used an RNAseq-based experimental approach. *X. birchmanni* and *X. malinche* individuals born to wild mothers (collected at the Coacuilco and Chicayotla populations respectively [64]) were raised at 22.5°C (14h light:10h dark cycle). A separate group of *X. malinche* females were crossed with *X. birchmanni* males to generate F_1_ hybrids. All individuals were raised in the same lab environment to adulthood before experiments began. Though we cannot discount the potential impact of maternal effects on expression response, all mothers were reared under the same environmental conditions.

For thermal stress experiments, three male individuals from each group were kept at a control temperature of 22.5°C for the duration of the experiment, and three male individuals from each group were subjected to a thermal stress treatment. Males in the treatment trials experienced a temperature ramp-up of 0.3°C/min from 22.5°C to 33.5°C (∼30 min duration). Control and treatment trials were run simultaneously between 11:00 AM and 1:00 PM. An air bubbler was used to maintain dissolved oxygen saturation in tank water for the duration of the experiment. Fish from both control and treatment tanks were anaesthetized with Tricaine mesylate diluted in tank water immediately after treatment tanks reached 33.5°C, and brain and liver tissues were dissected and placed in RNAlater. These samples were stored at 4°C for 24 hours and subsequently at -20°C. mRNA was extracted for a total of 36 brain and liver samples with a Qiagen RNeasy MiniPrep Kit. One *X. birchmanni* brain from the 22.5°C treatment and one *X. malinche* brain from the 33.5°C treatment yielded insufficient mRNA for RNAseq library preparation; therefore, these samples were not sequenced. RNAseq libraries were prepared using a KAPA mRNA HyperPrep Kit, pooled, and sequenced on three Illumina HiSeq4000 lanes. To control for batch effects, extraction, library prep, and sequencing batches were designed to include one individual from each biological group. We sequenced three biological replicates per experimental group and collected >30 million 150 bp paired-end reads per sample (Table S2).

### Differential gene expression analysis

Genes that are differentially expressed in response to thermal treatment, especially those that respond differently in *X. birchmanni* and *X. malinche*, are candidate genes that may contribute to variation in thermotolerance between species. For differential gene expression comparisons, we aligned RNAseq reads to reference transcriptomes inferred from high-quality *X. malinche* and *X. birchmanni* genome assemblies. For GO and KEGG enrichment analyses, we aligned reads to developed “pseudoreference” transcriptomes for these two species (from Schumer et al [77]) that were based on the genome assembly of the southern platyfish *X. maculatus* [78]. We used these references because *X. maculatus* is widely used as a model in melanoma research, and as a result has a well-annotated genome [78,79] with GO and KEGG pathways associated with each Ensembl gene ID. To reduce mapping bias in differential expression analysis we used a version of these references with within-species polymorphisms masked [77].

Before aligning reads, the program cutadapt and the FastQC wrapper tool Trim Galore! were used to trim Illumina adapter sequences and low-quality bases (Phred score < 30) from reads. All trimmed reads are available under NCBI BioProject PRJNA746324. One F_1_ liver sample from the 22.5°C ambient temperature treatment group was removed from downstream analyses due to unusually low read count (<1500 reads). Reads were then pseudoaligned to the *X. birchmanni* reference transcriptome with *kallisto* [80] and raw transcript counts were imported into the R package DESeq2 [81] for differential gene expression analysis. Genes with zero counts for all samples, extreme outliers (using a Cook’s distance cutoff of 0.99), or low mean normalized counts (i.e. genes with counts below an optimized threshold through an internal filtering step in DESeq2) were removed from analysis. This resulted in an analysis of 24,174 genes for both the brain and liver datasets.

To analyze differential expression of these genes, we used a design formula that included sequencing batch, genotype (*X. birchmanni, X. malinche*, or F_1_), and temperature treatment. Using DESeq2, we normalized gene counts by library size, estimated within-experimental group dispersion, fit a negative binomial generalized linear model, and tested significance with a Wald test. Following these steps, shrunken log-fold changes were calculated using an adaptive shrinkage estimator with a fitted mixture of normal distributions as a prior, derived from the ‘ashr’ package [82]. Genes were considered to be significantly differentially expressed between groups and treatments at an FDR-adjusted p-value < 0.1. To check for potential bias as a function of the reference sequence used in the pseudoalignment step, we repeated the above steps using the *X. malinche* reference transcriptome. Reassuringly, qualitatively similar results were obtained from this analysis (Supporting Information 2, Fig. S2).

### Co-expression network analysis with WGCNA

To identify clusters of interacting genes that respond to temperature treatment, we used the R package WGCNA to evaluate patterns of co-expression in the RNAseq data [83]. Weighted co-expression network analysis clusters genes with highly correlated expression patterns across samples into groups called modules. The expression patterns of modules are summarized by their ‘module eigengenes,’ defined as PC1 of the expression profiles of genes in the module, which can then be used to test for correlations between gene modules and traits or treatments of interest. This unsupervised approach is particularly powerful for identifying biological pathways whose expression strongly correlates with a specific treatment. In this case, we were most interested in modules that correlated with temperature treatment and with genotype.

WGCNA analysis was performed separately for sets of samples of each tissue type, using raw gene counts obtained from pseudoalignment to the *X. birchmanni* pseudoreference transcriptome (as described in the *Differential gene expression analysis* methods). We used the DESeq2 `varianceStabilizingTransformation` function to normalize raw gene counts by library size and size factors (the median ratio of the geometric mean of a gene over all samples) so that samples had comparable variances. Genes were filtered as described in the previous section, and additionally all genes with zero counts for one or more samples were dropped (out of 19,176 genes, 262 genes from the brain and 905 from the liver were dropped in this step).

As recommended by the WGCNA documentation, we selected a soft-thresholding power to transform the network into a more scale-free topology, which has been shown to better approximate biologically-relevant gene networks [84]. This step is intended to minimize the effect of noise in subsequent clustering steps and avoid using arbitrary thresholds for cluster construction. For each tissue dataset, the soft-thresholding power parameter was chosen by calculating the scale-free topology fit index for a range of powers (1 to 30) and selecting the asymptote of soft-thresholding power for downstream analysis (here, using the recommended thresholds for scale-free topology fitting index R^2^ > 0.8 and mean connectivity < 100). This resulted in the selection of soft-thresholding powers of 7 for the brain and 12 for the liver tissue datasets. Using these soft-thresholding values, we constructed single-block unsigned networks using WGCNA’s `blockwiseModules` function (see Appendix 1). We used a minimum module size of 20 and an unsigned topological overlap matrix to create a network that clusters genes by strength of co-expression, regardless of whether the correlation in expression is positive or negative.

After co-expression modules were identified using this approach, we sought to determine whether variation in any of these modules correlated with variation in traits of interest. As such, we looked for correlations between the module eigengene and genotype (*X. malinche, X. birchmanni, X. malinche-X. birchmanni* F_1_), temperature treatment (22.5°C and 33.5°C), or both. Correlations between traits and modules were calculated using the WGCNA `corPvalueStudent` function, and modules that correlated with genotype, temperature treatment, or both at Student asymptotic p-value < 0.05 were selected for further analysis (31 out of 54 for brain, 16 out of 50 for liver).

### GO and KEGG pathway enrichment of differentially expressed genes between temperature treatments

To explore which biological pathways are most affected by temperature treatment, we performed KEGG pathway and Gene Ontology (GO) term enrichment analysis of differentially expressed genes identified using results from the analyses of the brain and liver RNAseq data described above. We asked if there were enriched KEGG pathways in the set of genes that were significantly differentially expressed between *X. birchmanni, X. malinche*, and F_1_s at each temperature treatment (FDR adjusted p-value < 0.1). To do so, Gene IDs were mapped to Entrez IDs using the *X. maculatus* Ensembl database (version 99), and enriched KEGG pathways for each dataset were generated with the `kegg.gsets` function from the R package GAGE [85].

For GO enrichment, the R packages biomaRt [86] and GOstats [87] were used to extract *X. maculatus* Ensembl IDs and generate a GO gene universe of all genes analyzed with DESeq2, as described above (19,143 genes for brain, 19,176 for liver). We used a hypergeometric test implemented in the R package GSEABase [88] to identify overrepresented GO terms in the set of significantly differentially expressed genes between temperature treatments. We also performed GO analysis of genes in gene modules that correlated with temperature treatment in WGCNA analyses (12 modules for brain, 2 for liver). For both sets of GO analyses, we focused on significantly enriched categories (hypergeometric test p-value < 0.05) where greater than one gene was observed in our focal dataset.

### Ancestry of QTL and circadian clock genes in natural populations

We analyzed data from naturally occurring *X. malinche-X. birchmanni* hybrid populations to evaluate evidence for shifts in ancestry at genes under the chromosome 22 QTL, and clock genes that show misexpression in hybrids (see Results). These data from natural hybrid populations have been published in previous studies [72,77], so we only describe it briefly here. We analyzed data collected from the Tlatemaco (n=85) and Acuapa (n=97) hybrid populations in 2017 and 2018, respectively [72,77] (Fig. 1A). We previously collected ∼1X whole genome sequence data from individual hybrids collected from these populations and followed the local ancestry inference approach described above except that we used population-specific priors for admixture proportion and time since initial admixture. This approach resulted in estimates of the posterior probability for ancestry state at ∼613-629 million ancestry informative sites throughout the genome in the two populations. Using ancestry informative sites that fell within annotated coding regions, we generated estimates of the average ancestry per gene in both natural hybrid populations. We compared ancestry at genes of interest to the genomic background of each population.

### Correlation of caudal spot phenotype and CT_max_ in natural hybrids

Hybrids between *X. malinche* and *X. birchmanni* harbor a number of extreme traits not seen in either of the parental species. One of these is a hybrid incompatibility that causes melanoma in some individuals, originating from a melanocyte spotting pattern on the caudal fin [89]. To evaluate any relationship between this spotted caudal phenotype and CT_max_, we measured CT_max_ using the methods described above (see *Comparison of CT*_*max*_ *between X. malinche, X. birchmanni, F*_*1*_*s, and F*_*2*_*s and measurement of CT*_*max*_ *for QTL mapping*) for 123 lab-raised natural hybrids born from wild-caught mothers from the Chahuaco Falls population. These 123 natural hybrids were reared to adulthood under common conditions in the lab. Individuals were classified as having one of the following caudal spot phenotypes: no spot, normal spotted caudal, expanded spotted caudal, and 3D melanoma. Past histological work has indicated that individuals with the expanded spotted caudal phenotype have early-stage melanoma [89]. Individuals were assigned a 3D melanoma phenotype if they had melanoma that had completely overtaken the caudal fin (i.e. completely melanized and/or degrading) and/or that was growing laterally off the fin. We used a linear model to test for a correlation between CT_max_ and caudal spot phenotype.

## Results

### Evidence for a genetic basis for variation in thermotolerance

Given that *X. malinche* and *X. birchmanni* live in different thermal environments (Fig. 1C), we predicted that these species may have adapted to their respective thermal ranges. To determine whether there was a genetic basis for variation in CT_max_ between *X. malinche* and *X. birchmanni*, we reared *X. malinche, X. birchmanni*, F_1_, and F_2_ hybrid fish in a common garden, and measured their CT_max_ (see Methods). We found that *X. birchmanni* have significantly higher CT_max_ than *X. malinche* (p-value < 10^−6^; see Table S1), and that F_1_ and F_2_ hybrids exhibit intermediate CT_max_ (Fig. 2A). Though we know that CT_max_ is partially environmentally mediated in this system [65], this result shows that variation in CT_max_ between these species is also partly attributable to genetic differences.

**Figure 2.**
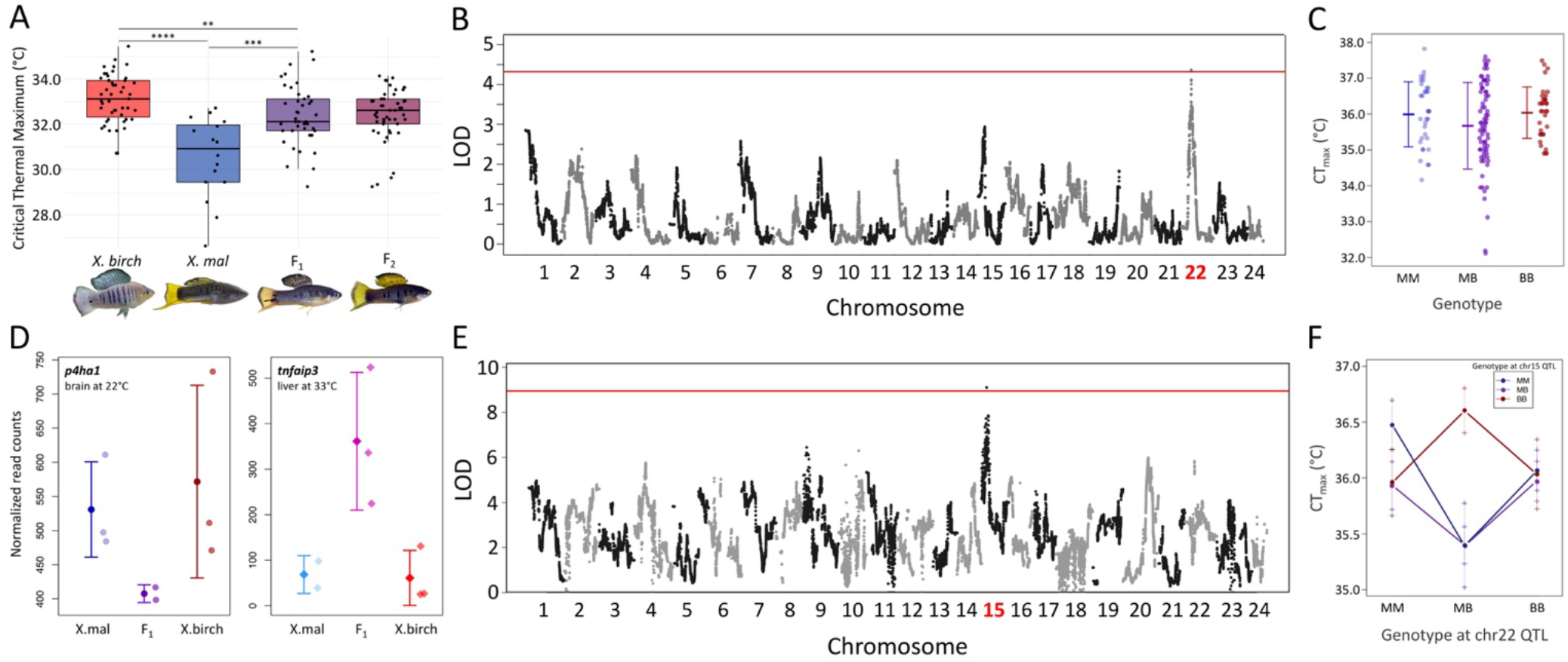
**A**. Results of CT_max_ trials on parental and hybrid individuals raised under common garden conditions indicate that variation in thermal tolerance between *X. birchmanni* and *X. malinche* is controlled in part by genetic factors. *X. birchmanni* has a significantly higher CT_max_ than *X. malinche*, and F_1_s and F_2_s on average have an intermediate CT_max_. See Table S1 for p-values for statistical comparisons between groups using Mann-Whitney Wilcoxon test. **B**. QTL mapping reveals one region on chromosome 22 associated with CT_max_. The QTL is significant at a 10% false discovery rate threshold, determined by permutations (red line). **C**. Artificial hybrids individuals with a heterozygous genotype at the peak associated marker on chromosome 22 have a 0.3°C reduction in CT_max_ on average compared to hybrid individuals homozygous for *X. malinche* or *X. birchmanni* ancestry, which have comparable CT_max_ on average. Bars and whiskers show the CT_max_ means for each genotype and 1 standard deviation. Points represent the CT_max_ of individual hybrids. **D**. Of the 45 genes under the CT_max_ QTL on chromosome 22, several show misexpression in F_1_s in at least one tissue or thermal context. The two examples shown here are *p4ha1*, which has significantly reduced expression in F_1_ brains at ambient temperature, and *tnfaip3*, which has significantly higher expression in F_1_ liver tissue under thermal stress (both at FDR adjusted p-value < 0.1). In these expression plots, mean normalized counts at 22.5°C are represented by a circle in a darker color and mean normalized counts at 33.5°C are represented by a diamond in a brighter color. Error bars show one standard deviation of expression. **E**. A second QTL scan, adding genotype at the chromosome 22 QTL as an interaction term, uncovered a putative interacting QTL on chromosome 15. This QTL is significant at a 10% false discovery rate threshold, determined by permutations (red line). **F**. Interaction plot of the peak associated marker of the chromosome 22 QTL (on the x-axis) and the peak associated marker of the chromosome 15 QTL (in the legend). This analysis shows that a combination of a heterozygous or homozygous *X. malinche* ancestry at the chromosome 15 QTL and a heterozygous genotype at the chromosome 22 QTL is associated with reduced CT_max_. By contrast combination of homozygous *X. birchmanni* ancestry and heterozygous genotype at the chromosome 15 and 22 QTLs, respectively, is associated with a modest increase in CT_max_ (see Table S5 for adjusted p-values). Bars and whiskers show the mean and 1 standard error.

### QTL mapping of loci involved in thermotolerance

Given these results, we proceeded to perform QTL mapping to evaluate associations between CT_max_ and ancestry in *X. malinche-X. birchmanni* artificial hybrids raised under common conditions (see Methods). We detected a single QTL associated with CT_max_ at a 10% false discovery rate threshold (Fig. 2B). The 1.5 LOD interval of this QTL spans ∼2.5 Mbs and contains 45 genes.

Surprisingly, further examination revealed that the QTL we detected was not associated with species-level differences in CT_max_. Instead, heterozygous ancestry in this region was associated with an average reduction in CT_max_ of 0.3°C (Fig. 2C). We estimate this QTL to have a moderate effect on the overall variation in CT_max_ in artificial hybrids, explaining approximately 6.9% of the variation (see Methods, Supporting Information 1, and Fig. S1 for simulations evaluating effect size inflation).

The relationship between genotype and phenotype observed at the chromosome 22 QTL is consistent with underdominance. Individuals with either homozygous genotype exhibit approximately the same average CT_max_ whereas individuals heterozygous for ancestry have a significantly reduced CT_max_ on average (Table S5; Fig. 2C). Though distinguishing whether this signal is caused by true underdominance or pseudo-underdominance (i.e. two or more linked loci with dominance and opposing effects in homozygotes; Fig. S3) is not feasible with our data, we discuss this possibility in more detail in Supporting Information 4.

### Exploring candidates in the QTL region

There are several possible explanations for the observed signal of underdominance at the chromosome 22 CT_max_ QTL. Because chromosomal inversions are a common genetic cause of underdominance [90–93], we confirmed that there are no inversions under this QTL by aligning *X. malinche* and *X. birchmanni* PacBio assemblies (Fig. S4; Supporting Information 4).

Next, we investigated genes that fell within the QTL region. The 1.5-LOD interval associated with the QTL spans from ∼8.6 Mb to 10.1 Mb, overlapping with 45 genes. We investigated functional annotations and patterns of expression of genes in this region, as well as amino acid differences between species (see Supporting Information 3). Because thermotolerance is a complex trait that is impacted by many biological pathways, narrowing down causal loci under the QTL based on their annotations is not straightforward. We highlight a handful of candidate genes in Table S3 but focus primarily on candidates with notable expression patterns in this section.

Given the CT_max_ phenotypes observed in individuals heterozygous for ancestry at this QTL, we were particularly interested in comparing gene expression patterns in F_1_ hybrids to the parental species. Because heterozygotes have reduced CT_max_ on average, we might expect genes controlled by a causal locus in *cis* to be misexpressed in F_1_ hybrids (which are heterozygous for ancestry across the whole genome). We evaluated expression patterns in the brain and liver and identified genes that were misexpressed in F_1_ hybrids compared to *X. birchmanni* and *X. malinche*, defined here as significantly higher or lower than either of the typical parental expression ranges, in either temperature condition. Of the genes in the QTL interval that were significantly differentially expressed between *X. malinche* and *X. birchmanni* for at least one of the temperature treatments (17 in the brain, 7 in the liver), most mirrored expression levels of one of the parental species or had intermediate expression in F_1_s (Table S3). However, four genes in this interval (*p4ha1, ndnf, tnfaip3*, and *infgr1l*) were misexpressed in F_1_s in at least one tissue and temperature condition (Fig. 2D, S8). We also identified genes that responded differently to thermal stress in F_1_s compared to parentals (see brain and liver expression results in Tables S6 and S7, respectively). For example, the zinc-finger protein *zfp62* exhibited an exaggerated response to high temperature compared to parental expression responses and the spliceosome subunit *sf3b5* was significantly downregulated at high temperatures in F_1_s whereas parental expression remained constant across temperatures (Fig. S8).

### Detection of a possible interacting QTL on chromosome 15

Another potential cause of a signal of underdominance at the chromosome 22 QTL could be interactions with other regions of the genome. In particular, in the literature on the evolution of gene regulation, a breakdown in interactions between paired *cis-* and *trans-*acting regulatory elements can explain aberrant expression patterns in F_1_ hybrids [36,43,94]. Thus, we performed a second QTL scan, including genotypes at the chromosome 22 QTL peak as an interaction term. Based on this analysis, we recovered a second QTL at the permuted 10% false discovery threshold, spanning ∼2.1 Mb on chromosome 15 (Fig. 2E).

While we are cautious of overinterpreting this result given low power in our study, we discuss it briefly here. Intriguingly, we find that artificial hybrids heterozygous at *both* the chromosome 22 and 15 QTL have reduced CT_max_ on average (−0.4°C), but that hybrids heterozygous at the chromosome 22 QTL and homozygous *X. birchmanni* at the chromosome 15 QTL have elevated CT_max_ on average (+0.5°C over hybrids homozygous for the *X. malinche* allele; Table S5; Fig. 2F, S5). We estimate that the combined additive and interaction effects of the chromosome 22 and 15 QTL could explain ∼14.8% of the total variation in CT_max_ in the artificial hybrids, although this number is likely an overestimate of their true effect size (see [76] and Supporting Information 1).

We explored evidence for known genetic interactions between genes under the chromosome 22 and chromosome 15 QTLs. The zinc finger protein gene *zbtb18* and the adjacent serine/threonine-protein kinase *akt3*, and the heterogenous nuclear RNA binding protein *hnrnpu*, which fall directly under the chromosome 22 and 15 peaks respectively, are known to interact during neurodevelopment [95]. We discuss what is known about their interactions and other interacting genes in these regions in more detail in Supporting Information 5 and summarize all genes under the chromosome 15 QTL in Table S4.

### Gene expression profiles differ between species and thermal treatment

To broadly survey changes in gene expression between parental species and their F_1_ hybrids in response to high temperature, we generated RNAseq data for brain and liver tissue from fish exposed to ambient and high temperature conditions (see Methods). We sampled the brain and liver to survey two tissues that play different roles in organismal homeostasis – energy consumption and detection of and response to environmental changes by the brain, and energy metabolism by the liver.

In addition to using these data to evaluate expression patterns of genes under the QTL region, we analyzed it in a genome-wide context. As expected, we found broad differences in expression between tissues and species (Fig. S6), as well as strong responses to high temperature (Fig. 3A, S7). The vast majority of the variation in expression in our dataset is explained by tissue type (83.5%; Fig. S6); therefore, the tissue datasets were analyzed separately. Of the remaining variation, genotype explained ∼23% of the variation in expression between samples and temperature treatment explained ∼11% (Fig. 3A). A large number of genes were significantly differentially expressed between *X. birchmanni* and *X. malinche* (FDR adjusted p-value < 0.1) across temperatures in both tissues (brain: 3,357 and 3,121 genes; liver: 2,318 and 1,508 genes at 22.5°C and 33.5°C, respectively). Interestingly, while the number of genes for which expression changed in response to high temperature in *X. birchmanni* and *X. malinche* was similar (brain: 882 and 979 genes; liver: 113 and 38 genes, respectively), ∼2.5-3.5x more genes were responsive to high temperature in F_1_ brain (2,318) and liver (408) tissues compared to those of parentals. We found that ∼3% and ∼0.5% of all genes across the genome were misexpressed under at least one of the two tested thermal contexts in F_1_ hybrid brains and livers, respectively. Moreover, ∼9% and ∼3% of the genes that responded to temperature in *X. birchmanni* and/or *X. malinche* were misexpressed under at least one of the thermal contexts in F_1_ brains and livers, respectively. Overall, we found that more genes were misexpressed (low or high) in F_1_ hybrids at 22.5°C than at 33.5°C in the brain (521 versus 187, respectively), whereas the liver exhibited the opposite pattern (10 versus 96, respectively); however, we are cautious of potential technical factors that could influence these patterns, such as differences in variance between groups at high temperature. Only a handful of genes were misexpressed under both thermal contexts. See Tables S6-7 for complete results for each group, and Table S8 for a summary of F_1_ expression patterns.

**Figure 3.**
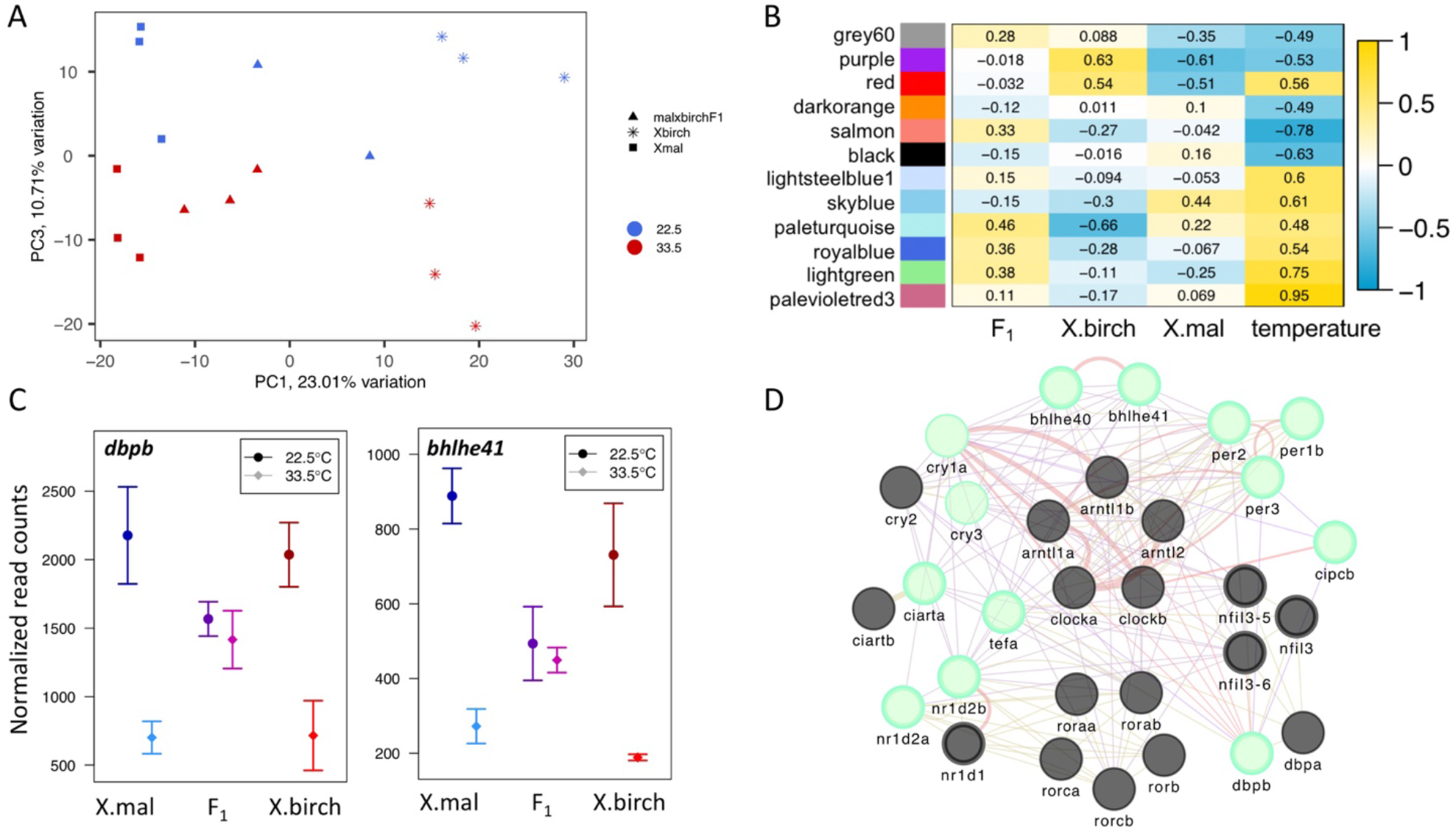
**A**. PCA plot of normalized gene count data in the brain for all 17 individuals for which RNAseq data was collected. Individuals clearly separate by genotype and temperature treatment along PC1 and PC3 respectively. Genotype explained 23.01% of the variation in overall expression and temperature treatment explained 10.71%. PC2, which is not shown here, explained 19.22% of the variation in expression and was most strongly correlated with sequencing batch. **B**. Weighted gene co-expression analysis uncovered 12 temperature-associated modules in the brain (shown here) and 2 in the liver (Fig. S9). Traits are listed on the x-axis, and color blocks and labels on the y-axis represent the WGCNA module. Pearson’s correlation coefficients are listed for each module and trait, with box color corresponding to the strength of the correlation (yellow spectrum for a positive trait-module correlation, blue spectrum for a negative trait-module correlation). **C**. Several clock genes that were identified in the circadian clock gene-enriched module, including *dbpb* and *bhlhe41*, are misexpressed under both ambient and high temperature conditions in F_1_ brains. Interestingly, the mechanism of misexpression may be due to a failure of F_1_ hybrids to respond to temperature change. *X. birchmanni* and *X. malinche* strongly downregulate both genes in response to high temperature, while F_1_s do not. **D**. The gene network for core circadian clock genes in the *Xiphophorus* genome, predicted by GeneMania [128] and visualized with Cytoscape [129]. The structure of the network is colored based on the nature of evidence of each interaction, including direct interactions between genes (red), co-expression (purple), and shared domains (yellow). Genes that are misexpressed in F_1_ brains at high temperature in our study are highlighted in bright green, and genes that appeared in the circadian clock gene expression module identified by WGCNA are shown with a bold outline.

We report functional categories with enriched expression responses (p-value < 0.05) to thermal stress, including general thermal stimulus and immune response pathways, in Table S9. In the brain, 84 GO terms are enriched in response to temperature in both *X. malinche* and *X. birchmanni*. Notably, response to temperature stimulus and circadian rhythm categories were commonly enriched for both species. Additionally, 77 terms are enriched only in *X. birchmanni*, and 70 terms are enriched just for *X. malinche*. Among those terms that were enriched only in *X. birchmanni* under thermal stress were autophagy and disassembly of mitochondria, negative regulation of biosynthesis and gene expression, and endogenous stimulus response pathways.

KEGG analysis recovered only one biological pathway commonly enriched in the set of genes that were significantly differentially expressed between temperature treatments in *X. malinche* and *X. birchmanni* brains: protein processing in the endoplasmic reticulum (xma04141; FDR adjusted p-value < 0.1; Table S10). This result may be attributable to the fact that the endoplasmic reticulum plays a key role in the unfolded protein response, which is activated by thermal stress and is key for maintaining homeostasis during stress [96]. Intriguingly, one transcriptional activator of the unfolded protein response, *xbp1*, is significantly upregulated in both *X. malinche* and *X. birchmanni*, but not in F_1_ hybrid brains (Fig. S8).

Strikingly, 20 KEGG pathways were significantly enriched in F_1_ brains in response to high temperature. Among these enriched pathways were protein processing in the endoplasmic reticulum and signaling pathways (see Table S10 for full list) that induce the transcriptional regulator of the innate immune response, *nfkb1* [97–100]. Interestingly, one of the potential sets of interactors under the chromosome 22 and chromosome 15 QTL are inhibitors of *nfkb1* expression (see Supporting Information 5 for more information). No significantly enriched KEGG pathways were recovered from the set of genes differentially expressed in response to temperature in the liver across groups.

### Co-expression network analysis reveals misexpression in F_1_ clock genes

We used the co-expression network analysis software WGCNA [83] to identify clusters of co-expressed genes in our RNAseq datasets (Fig. S10, S11). We performed this analysis separately for the two tissue types. In total, 54 and 50 gene co-expression modules were recovered from the brain and liver RNAseq data, 12 and 2 of which were significantly correlated with temperature treatment respectively (p-value < 0.05; Table S11; Fig. 3B, S9). Additionally, four of the 12 brain temperature-correlated modules were significantly correlated with at least one genotype (see Supporting Information 8).

Notably, one temperature-correlated module was shared between tissue types, suggesting that it may represent a cluster of genes globally involved in the thermal stress response. This module is enriched in genes involved in the circadian rhythm and circadian regulation. This finding is notable since circadian clock pathways are impacted by temperature and play a role in thermoregulation and thermal stress response across taxonomic groups [101–104].

Strikingly, several of the circadian clock genes in this shared temperature-associated module are misexpressed in F_1_ hybrids, particularly in data collected from the brain in the high temperature treatment (Supporting Information 6-7). The number of misexpressed genes in this module greatly exceeds the number expected by chance (based on permutations, Table S12, Fig. 3D, Supporting Information 7). This suggests that genes in these circadian clock pathways may be commonly misregulated under thermal stress in *X. malinche-X. birchmanni* hybrids. Specifically, we find that most of the clock genes in this module are strongly up- or down-regulated in *X. malinche* and *X. birchmanni* brains and livers in response to high temperature. In contrast, at ambient temperature, F_1_ clock gene expression tends to be similar to parental expression, but at high temperatures these genes are misexpressed in F_1_ brains compared to parental brains (Table S13; Fig. 3C-D). These results hint at a failure to regulate expression of these genes in hybrids. Specifically, much of the misexpression observed in these genes is attributable to the fact that while their expression in parental brains is strongly responsive to the thermal treatment, F_1_ expression does not change substantially between temperature treatments. Additionally, some of these genes, such as the transcription factors *dbpb* and *bhlhe41* shown in Fig. 3C and *nr1d2a* and *cipcb* shown in Fig. S8, show patterns of F_1_ misregulation under both thermal contexts. We discuss these patterns in more detail in Supporting Information 9.

### Ancestry patterns in natural hybrid populations at regions implicated in thermotolerance

Hybrid populations between *X. birchmanni* and *X. malinche* occur across a range of elevations in different river systems [64] and experience different average temperatures [65]. To determine whether there is evidence of selection against a particular ancestry state in natural hybrid populations at the chromosome 22 and 15 CT_max_ QTL and at the clock genes discussed above, we focused on two hybrid populations that occur at elevations closer to those typical of *X. birchmanni* populations (Fig. 1A) and thus experience higher temperatures on average. These populations are the Acuapa and Tlatemaco populations (elevations of 476 and 480 meters, respectively). Notably, while individuals from the Acuapa population derive the majority of their genomes from *X. birchmanni* (∼75%; [72]), the parental species with higher thermotolerance, individuals from the Tlatemaco population derive the majority of their genomes from *X. malinche* (∼72%; [67]). Thus, regions that have unusually high *X. birchmanni* ancestry in both populations compared with the genome-wide background and that overlap with mapping or expression results may be of particular interest as candidates for loci underlying variation in thermotolerance phenotypes.

Focusing first on the QTL regions, we found that four genes under the chromosome 22 QTL (*akt3, sdccag8*, and *olig3*) and a handful of genes under the chromosome 15 QTL (including *nrxn3a* and one *nrxn3b* isoform) have higher than average *X. birchmanni* ancestry in both low-elevation hybrid populations (Fig. 4A). This shared high *X. birchmanni* ancestry in both populations deviates significantly from expectation (based on permutations, Table S14).

**Figure 4.**
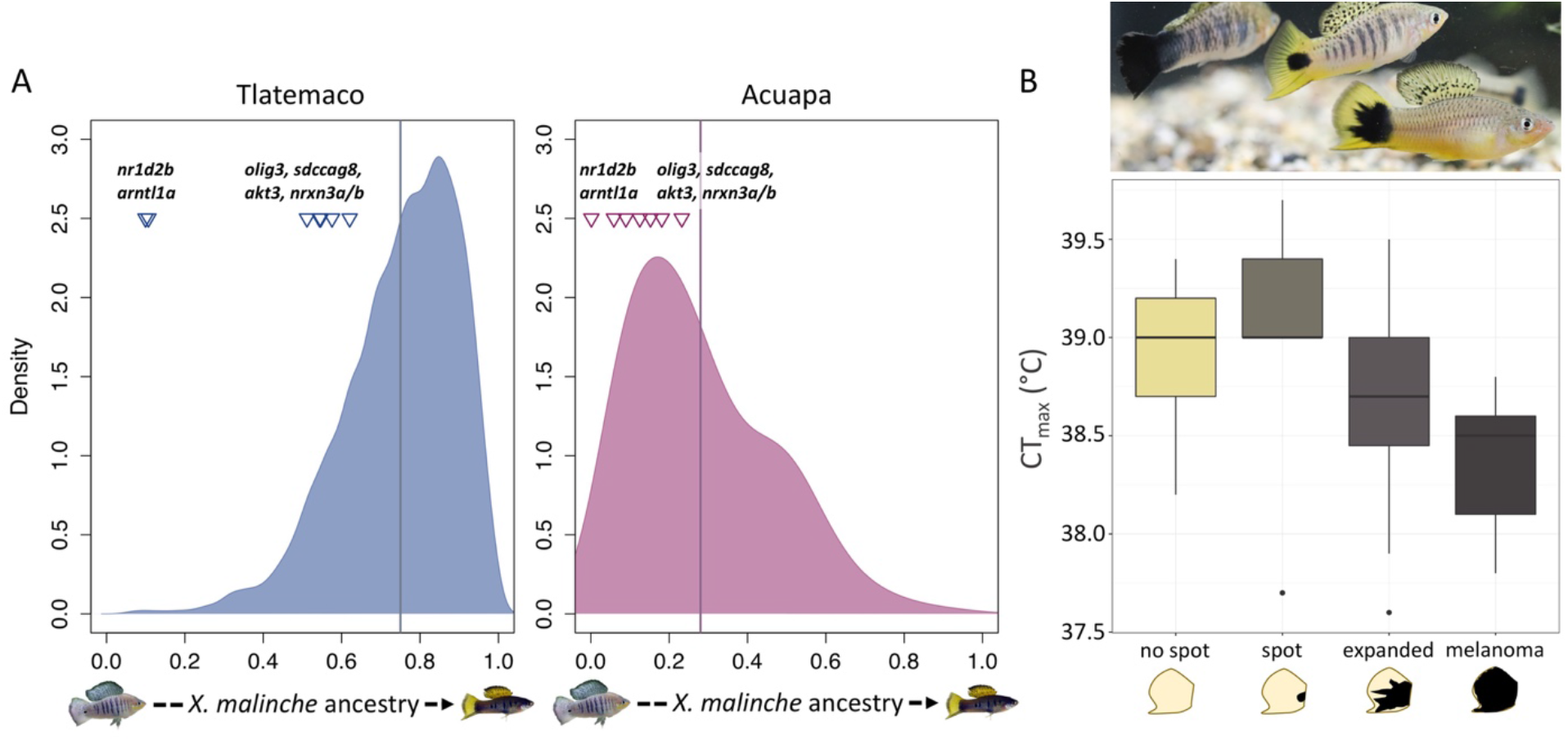
**A**. Ancestry at regions implicated in thermal stress response compared to the genome-wide ancestry distributions in two natural hybrid populations that occur at low elevations. Individuals from the Tlatemaco population derive on average ∼75% of their genome from the *X. malinche* parent species and individuals from the Acuapa population derive on average ∼28% of their genome from the *X. malinche* parent (genome-wide means represented by solid lines). Conversely, a handful of genes under the chromosome 22 (*olig3, sdccga8, akt3*) and 15 (*nrxn3a, nrxn3b*) QTL and two clock genes (*nr1d2b, arntl1a*) have unusually high *X. birchmanni* ancestry in both populations, raising the possibility that there may be positive selection for *X. birchmanni* ancestry at these genes in low elevation populations (see Table S14 for p-values from permutations). **B**. The top image shows three Chahuaco Falls hybrids, from left to right, with 3D melanoma, normal spotted caudal, and expanded spot phenotypes. Boxplots show CT_max_ of lab-reared Chahuaco Falls hybrids, split by spotted caudal phenotype. Lab-reared individuals with expanded spot and 3D melanoma phenotypes have significantly lower CT_max_ compared to individuals with no spot or a normal spotting pattern.

We next evaluated ancestry in both hybrid populations among genes in the circadian clock gene expression module. Notably, two clock genes in this module that are misexpressed in F_1_ hybrids, *nr1d2b* and *arntl1a*, have unusually high *X. birchmanni* ancestry in both populations (>89% in both, permuted p-value<0.01; Fig. 4A, Table S14). Interestingly, *nr1d2b* directly represses *arntl1a* expression [105]. Clock genes with strong skews in ancestry in both natural hybrid populations may be adaptive in lower elevation habitats, as this level of ancestry sharing across the two populations is unexpected by chance (see Supporting Information 10). Together, these analyses highlight regions that may be under selection due to their impacts on thermotolerance in natural hybrid populations.

### Other phenotypes associated with thermotolerance in hybrids

Given the overall pattern of reduced thermotolerance associated with heterozygous ancestry at the chromosome 22 QTL and aberrant expression of many thermally responsive genes in F_1_ hybrids, we wanted to further investigate other possible phenotypic drivers of reduced thermotolerance in hybrids. One trait that is present in hybrids but not in parental individuals of either species is a hybrid incompatibility involving a pigmentation phenotype called the “spotted caudal”. While the spotted caudal is a benign melanocyte pigmentation pattern in *X. birchmanni*, it can transform into a malignant melanoma in hybrids (Fig. 4B) with certain genotype combinations ([89]; those with *X. birchmanni* ancestry at the *xmrk* gene and *X. malinche* ancestry at *cd97*).

We found that the spotted caudal phenotype was significantly correlated with CT_max_ in lab-reared offspring from wild mothers collected from a natural *X. malinche-X. birchmanni* hybrid population from Chahuaco Falls (Fig. 1A). In particular, hybrid individuals with an expanded spot typical of early melanoma as well as hybrids with a more advanced 3D melanoma phenotype had significantly reduced CT_max_ compared to those with a benign spot or no spot (Fig. 4B). This poor performance in hybrids with incompatible genotype combinations highlights one potential mechanism through which underdominance in traits such as thermotolerance could occur. We discuss the implications of this result in more detail in Supporting Information 11.

## Discussion

How adaptive traits arise at the genetic level has been a classic question in evolutionary biology for decades. Here, we used a QTL mapping approach to identify loci contributing to variation in thermotolerance in hybrids between the northern swordtail species *X. malinche* and *X. birchmanni*. Mapping CT_max_ QTL in an artificial hybrid population revealed one underdominant QTL spanning ∼1.5 Mb on chromosome 22 and a putative interacting QTL on chromosome 15. This finding, along with our gene expression results, points to a breakdown in the response to thermal stress in hybrids, with important implications for understanding the genetic architecture and evolution of ecologically relevant traits in general.

Though more commonly reported in plants ([90,106–108]; but see [109]), underdominant QTL provide insight into genotypes that may be disadvantageous in hybrids. For example, mapping pollen fertility in *Mimulus* has identified hybrid sterility loci in heterozygotes caused by structural rearrangements [90] and mapping in tomatoes has revealed a reduction in fruit size in heterozygotes [108]. Unlike these QTL, which generally appear to have a simple genetic architecture, we find that the QTL on chromosome 22 explains a modest proportion of the total variation in this trait in *X. malinche*-*X. birchmanni* artificial hybrids. This both highlights the complex nature of this trait, and explains why, despite an average signal of reduced CT_max_ in individuals heterozygous at the chromosome 22 QTL (Fig. 2C), most F_1_ and F_2_ hybrids have a CT_max_ that is intermediate to the parental ranges (Fig. 2A; Supporting Information 4).

What mechanisms drive reduced thermotolerance of heterozygous individuals at the chromosome 22 QTL? One clue comes from gene expression results from *X. malinche, X. birchmanni*, and F_1_ hybrids. We see widespread misexpression in F_1_ hybrids (approximately 9% and 3% of temperature responsive genes in parental brain and liver, respectively), where heterozygous individuals show expression patterns far outside the range of either parental species, including at genes under the chromosome 22 QTL (Fig. 2D). These aberrant expression patterns likely result from disruption of gene expression networks in hybrids at the molecular level [43], and could lead to phenotypic effects such as the reduced CT_max_ we observe at the chromosome 22 QTL. While well-documented in literature on the evolution of gene regulation [36,94,110–112], these types of misexpression dynamics have only recently been appreciated in the speciation genetics community as a source of hybrid incompatibilities between species [94].

One particularly intriguing example of gene expression misregulation in F_1_ hybrids occurs in circadian clock pathways. Overall, we find strong correlations between co-expression patterns of clock genes and temperature treatment in our RNAseq datasets. This finding is consistent with decades of data showing that expression levels of core clock genes are regulated in response to temperature across taxa (for example in plants: [104,113]; flies: [114]; fish: [115–119]; mammals: [120]). This regulatory response is important for maintaining homeostasis and timing of the biological clock regardless of temperature-induced shifts in basic processes like enzymatic activity [121]. While we observe a strong circadian clock regulatory response to temperature treatment in both *X. malinche* and *X. birchmanni*, we find that an unexpectedly large number of circadian clock genes are misexpressed in F_1_ hybrids (permuted p-value<10^−6^), particularly after exposure to high temperature (Fig. 3C-D; Supporting Information 7). The response observed in parent species suggests that proper regulation of these genes is important in thermal stress response in *Xiphophorus*, and enriched misexpression in F_1_ hybrids points to a potential breakdown of basic regulatory processes in hybrids. Moreover, multiple pairs of genes that fall under the chromosome 22 and 15 QTLs are known to interact with clock genes. For example, several loci under the QTL regions (*akt3, zbtb18, nrxn3b, tnfaip3*, and *nfkbia*) are co-expressed or interact with the regulatory clock gene *bhlhe40* [122–126]. Future work should address the functional basis of this misregulation as well as whether hybrids exhibit difficulty maintaining homeostasis compared to the two parental species, particularly at a range of rearing temperatures.

Consistent with a role in fitness in natural populations, we see evidence of selection on ancestry at a handful of temperature-associated clock genes. Natural hybrids from the Acuapa and Tlatemaco populations derive the majority of their genomes from *X. birchmanni* and *X. malinche*, respectively, but both reside at *X. birchmanni* typical elevations. Specifically, clock genes *nr1d2b* and *arntl1a* (Fig. 4A) are unusually skewed towards *X. birchmanni* ancestry in both populations. This could indicate an ecological advantage of the *X. birchmanni* alleles at these genes (or selection to resolve misexpression).

Given evidence for poorer performance and widespread misexpression in some hybrid individuals in response to thermal stress, we were curious about the ways that known hybrid incompatibilities interact with the thermal environment. Previous work has shown that in *X. malinche-X. birchmanni* hybrids, the combination of *X. malinche* ancestry at the gene *cd97* and *X. birchmanni* ancestry at the gene *xmrk* results in the formation of a malignant melanoma. This incompatibility appears to reduce fitness in the wild based on population resampling results, but the mechanism is unclear, as individuals can survive for more than 2 years in the lab even with severe melanoma [89]. We found that both 3D melanoma and less severe melanoma are significantly correlated with reduced CT_max_ in *X. malinche-X. birchmanni* hybrids. This hints at a potential ecological fitness consequence for individuals with the melanoma incompatibility and exploring whether this relationship is causal is an exciting future direction (we discuss this result more thoroughly in Supporting Information 11).

We set out to use QTL mapping and differential gene expression analysis to identify the genetic basis of differences in thermotolerance between *X. malinche* and *X. birchmanni*, so that we could identify regions of the genome that may undergo adaptive introgression in response to changing thermal environments. However, our mapping and RNAseq results instead uncovered signals of hybrid breakdown and potential BDMIs. Our results highlight a more general problem with QTL mapping of species-level differences; in some cases, breakdown in the biological processes and traits of interest in hybrids will obscure the differences between the parental species that researchers seek to map. On the other hand, our results provide indirect clues into the expected outcomes for our original questions. Hybrids between *X. malinche* and *X. birchmanni* experience widespread misregulation of genes that respond to thermal treatments in the parental species, and some individuals that harbor heterozygous ancestry at the chromosome 22 QTL or a common hybrid incompatibility between species exhibit markedly reduced thermotolerance. These results suggest that adaptive introgression of as of yet unidentified *X. birchmanni* thermotolerance alleles may not be sufficient to offset the costs of hybridization, and therefore may not lead to higher thermotolerance in *X. malinche* populations. We also note that although we focus on CT_max_ in the present study, *X. malinche* is found in environments with lower temperatures than those experienced by any other *Xiphophorus* species. Studying the genetic architecture of tolerance of cool temperatures in *X. malinche* may provide insight into the pressures driving regulatory divergence between species and misexpression in hybrids.

Together, this work highlights the potential for ecological incompatibilities to play a role in selection on *X. malinche-X. birchmanni* hybrids [46]. Nearly a decade of work has uncovered evidence for genetic incompatibilities between these two species, but most cases that have been evaluated in detail have focused on intrinsic hybrid incompatibilities [89,127]. Our results underscore how shifts in global climate may impact a suite of biological processes and exacerbate or uncover ecological incompatibilities in hybrids. Such potential consequences may limit the success of genetic rescue as an effective strategy for population conservation.

## Acknowledgements

We thank Lisa Couper, Pablo Delclos, Justin Havird, Erik Iverson, Katya Mack, and members of the Schumer and Rosenthal labs for helpful feedback and discussion. We are grateful to the Mexican federal government for permission to collect samples. We thank Stanford University and the Stanford Research Computing Center for providing computational support for this project. This work was supported by NIGMS Center of the National Institutes of Health under award number T32GM007276, a Human Frontiers in Science Program grant to MS (RGY0081), a SICB GIAR to CP, and an NSF GRFP to VG.

