## Supporting Information for "Genomic insights into variation in thermotolerance between hybridizing swordtail fishes"

*Supporting Information 1. QTL effect size estimates and simulations evaluating inflation*

We used the R/qtl function `fitqtl` to estimate the effect size of each mapped QTL on the observed variation in $CT_{max}$ in artificial hybrids. Based on this analysis, we estimated that the chromosome 22 QTL to explains 6.9% of the observed variation in $CT_{max}$, the chromosome 15 QTL explains 5.3%, and the interaction between these QTL explains 2.6% of the variation. Together, we estimate that the additive effects of the chromosome 22 and 15 QTL and their interaction explains ~15% of the variance in $CT_{max}$ observed in our artificial hybrid mapping population.

While we are able to directly estimate the effect size of each QTL region, we expect that these effect sizes are inflated. This is because we expect our experiment to have low statistical power given the number of artificial hybrids tested, and inflation in estimated effect sizes in QTL studies with low statistical power has been well documented [1]. This phenomenon, known as the Beavis effect or the Winner's curse, is the product of the fact that cases where noise in the data happens to go in the same direction as the signal in the data are more likely to be detected when true power to detect the QTL is marginal. This is especially a concern in cases such as ours where the QTL peak is close to the genome-wide significance threshold.

Thus, to explore the range of effect sizes that may be consistent with the QTL signal that we detect on chromosome 22, we used a simulation-based approach. Specifically, we used an approximate Bayesian computation (ABC) approach to interrogate the range of effect sizes consistent with the observed data. We recorded the average difference in $CT_{max}$ between the parental species and coded this as the variable $CT_{diff}$. We extracted observed genotypes and covariates at the QTL peak and used the following steps in ABC simulations:

1. We drew a random effect size ($chr22_{eff}$) from a uniform distribution ranging from 0-30% for the simulation.
2. For each individual, we determined their genotype at the QTL peak.
    a. For individuals with homozygous *X. birchmanni* or homozygous *X. malinche* genotypes, we generated their phenotype by drawing from a random normal distribution with the mean equal to the observed average $CT_{max}$ in the artificial hybrid population, and the variance equal to observed variance in $CT_{max}$ in the hybrid population.
    b. For individuals that were heterozygous for *X. birchmanni* and *X. malinche* ancestry, we performed the step described in *a* to determine a base $CT_{max}$ phenotype. We then calculated that individual's phenotype as $CT_{max}$ – $chr22_{eff}*CT_{diff}$.
3. Using these simulated phenotypes and observed genotypes and covariates, we performed a linear regression as we had for the real data.
4. We accepted or rejected the simulation based on the estimated effect size, with a 5% tolerance threshold.
5. For each accepted simulation we recorded the true effect size simulated.
6. We repeated this procedure until 500 simulations had been accepted.

We recovered a well-resolved posterior distribution of effect sizes based on these simulations. We were most interested in the range of effect sizes consistent with our data. Based on the posterior distribution, we estimate the 95% credibility intervals to be 2.5-20%, compared to the point estimate of 6.9% from analysis of the real data (Fig. S1).

*Supporting Information 2. RNAseq analysis using the X. malinche pseudoreference*

To evaluate the impact of the reference transcriptome used for pseudoalignment on downstream expression results, we repeated all analyses using the *X. malinche* reference transcriptome. We then compared the list of genes with significantly different expression identified by both analyses, and the estimated log-fold changes in expression of these genes. We pseudoaligned all RNA-seq reads to the *X. malinche* reference transcriptome with *kallisto* and ran a differential gene expression analysis with DESeq2 (as was performed with the *X. birchmanni* reference, see Methods). We subset the expression results to only include genes with one-to-one annotations between the two references. For each expression comparison between groups, we identified genes with significantly different expression (adjusted p-value < 0.1). We found that >90% of genes in each comparison (genotype, temperature) were identified as differentially expressed regardless of choice of reference transcriptome. See Table S15 for a summary of these results. In addition, we measured the correlation in estimated log-fold change among these genes and found high concordance in these estimates regardless of the choice of reference transcriptome (>0.95 adjusted $R^2$ for all comparisons; Fig. S2, Table S15).

*Supporting Information 3. Amino acid differences between species in genes under the chromosome 22 QTL*

In addition to using RNAseq data to identify expression differences between species in the genes under the chromosome 22 and 15 QTL, we sought to identify genes that had amino acid substitutions between *X. malinche* and *X. birchmanni.* To this end, we extracted predicted cDNA sequences for each gene from the *X. birchmanni* and *X. malinche* reference genomes and used the codeml program from PAML [2] to infer relative rates of synonymous and nonsynonymous substitutions (dN/dS) for all 45 genes in the 1.5-LOD interval (Table S3). Three of the 45 genes had a dN/dS that exceeded 1. However, in all three cases this signal was driven by an absence of synonymous mutations, and the number of nonsynonymous substitutions between species was not unusual compared to genes of similar length (based on dN calculated for genes genome-wide within 10% of their length; the genes for *sparcl1*, *LOC111606527,* and *fam149b1* fell in the 31%, 35%, and 65% quantiles of dN for genes of similar lengths, respectively).

We also evaluated whether there were differences in the rate of evolution on the *X. malinche* branch for a handful of candidate genes (Table S16) with many nonsynonymous mutations. Specifically, we used the codeml function implemented in PAML to compare the likelihood ratio of a model with a single omega, to a model where omega differed along the lineage leading to *X. malinche*. We specified *X. malinche* as the species of interest for detecting changes in rate because *X. malinche* lives in a high-elevation habitat ecologically distinct from other species in the genus [3]. We took advantage of genomes from 14 *Xiphophorus* species to increase the power of these tests. However, we found no evidence for accelerated evolution of these genes along the *X. malinche* lineage (Table S16).

*Supporting Information 4. Signal of underdominance at the chromosome 22 QTL*

In the main text, we refer to the QTL on chromosome 22 as "underdominant" because individuals heterozygous for *X. birchmanni* and *X. malinche* ancestry in this region have on average lower thermotolerance than homozygotes of either species. However, this signal is complicated by a number of factors. It is worth noting that it is not possible with our data to disentangle true underdominance generated by the effects of a single-locus and pseudo-underdominance generated by closely linked genes [4,5]. Pseudo-overdominance, where linked genes cause a phenotype that exceeds that of both parental species, has historically been explored as an explanation for QTL contributing to heterosis [4,6,7]. Though sparsely documented in the literature, pseudo-underdominant QTL could arise from the same principles, where alleles of two additive loci are tightly linked, such that in crosses individuals with heterozygous genotypes at both loci appear to have extreme phenotypes due to dominance (see Fig. S3 for a schematic representation; [4,8,9]).

Most of the known examples of pseudo-over or underdominance come from research in crops. In one QTL mapping study of cultivated tomatoes, researchers mapped total soluble solid yield and identified an overdominant QTL. Fine-mapping of this QTL revealed that two linked QTL actually drove the observed patterns [10]. The signal that we document on chromosome 22 could plausibly be driven by linked loci with opposing effects on thermotolerance. Given the difficulties of producing artificial crosses between *X. malinche* and *X. birchmanni* and conducting large scale thermotolerance trials, fine mapping the chromosome 22 QTL is not currently feasible but would be an interesting direction for future work.

To rule out structural rearrangements as an explanation for the signal of underdominance observed at the chromosome 22 QTL, we aligned *X. malinche* and *X. birchmanni* PacBio assemblies of chromosome 22. Briefly, we used the 'nucmer' program from MUMmer4 [11] with default parameters to generate coordinate and delta files. We found no evidence of structural differences along chromosome 22 between *X. malinche* and *X. birchmanni* (Fig. S4).

*Supporting Information 5. Potential interacting genes between the chromosome 22 and 15 QTL*

In a two-dimensional scan, we identify a locus on chromosome 15 (at an FDR threshold of 10%) that may interact with the QTL we initially identified on chromosome 22 (see main text). We were curious about whether there were known molecular links between the genes in these two regions. Review of the literature as well as gene network prediction with GeneMania [12] revealed several potential interactions between the chromosome 22 and putative interacting chromosome 15 QTL. As discussed in the main text, the neurodevelopmental genes *akt3*, *zbtb18,* and *hnrnpu* interact and are implicated in mammalian neurological diseases [13]. For example, haploinsufficiency and dysfunction of these three genes have been shown to be the primary drivers of the various detrimental phenotypes characterizing 1q43q44 microdeletion syndrome [13]. *zbtb18* is also known to directly interact with the neuronal cell surface protein *nrxn3b*, which also falls under the chromosome 15 peak [14,15]. However, we note that none of these genes have been shown to play a direct role in thermotolerance (though all four have several genetic or physical interactions with circadian clock genes [14,16], a temperature-sensitive biological pathway that is perturbed in $F_1$ hybrids).

We highlight two additional pairs of interactors here. *sf3b5* under the chromosome 22 QTL and *sf3b6* under the chromosome 15 QTL are physically interacting subunits of the SF3B complex of the U2-dependent spliceosome, which is known to be sensitive to temperature [17]. Moreover, the subunit *sf3b1* has been shown to regulate the transcription factor *hsf1* during the heat shock response in *C. elegans* [17]. Notably, *sf3b3, sf3b4*, and *sf3b5*, as well as several other members of the SF3B gene network (*phfa5a, snrpa1, snrpd1, snrpd2, snrpd3*, and *lsm8*), are significantly downregulated in $F_1$ hybrids at high temperatures.

Finally, the tumor suppressors, *tnfaip3* and *nfkbia* under the chromosome 22 and 15 QTL, respectively, both negatively regulate *nfkb1,* a major transcription factor in immune and inflammatory responses [18]. Constitutive downregulation of these genes can result in excess inflammation and facilitate progression of various inflammatory diseases [19]; however, proper activation of *nfkb1* in response to high temperature has been shown to minimize heat stress-induced apoptosis [20]. Intriguingly, both genes show misexpression, or extreme expression compared to parental expression, in $F_1$ livers at the high temperature treatment (Fig. 2D, S8), suggestive of regulatory breakdown in the inhibitory pathway of *nfkb1* (though the same pattern is not observed in brain tissue). Additionally, both genes are strongly downregulated in *X. birchmanni* livers, but not in *X. malinche* livers, in response to high temperature, hinting at a potential regulatory mismatch in hybrids. However, we note that we do not observe aberrant expression of *nfkb1* at either temperature treatment in $F_1$ hybrids.

*Supporting Information 6. Constitutive misexpression of genes with high connectivity*

One of the signals that we detect in our dataset is widespread misregulation of genes in $F_1$ hybrids. In many cases, this misregulation may not specifically be related to thermotolerance phenotypes but is still of interest in understanding gene regulation in hybrids. We might expect that misregulation of hub genes, or genes that are highly connected and co-expressed with many other genes, would be more likely to have substantial consequences on gene network stability and drive downstream fitness consequences. We also might expect hub genes to be misregulated more frequently from first principles. Because hub genes have more genetic interactions by definition, they may be more likely to be misregulated in $F_1$s, as there are more partners modulating them. To explore whether hub genes are more or less likely to be misexpressed, we compared the number of observed misexpressed hub genes to expectations from permuted null distributions.

Hub genes were defined using scores for gene connectivity and gene module membership based on WGCNA analysis. First, we identified genes whose expression was strongly correlated with at least one module's eigengene (called the module membership; [21]). An arbitrary module membership cutoff of 0.85 was used as an initial filter (following [22]). Two measures of connectivity calculated by WGCNA were used to identify two focal sets of hub genes: within module (kWithin) and total (kTotal) connectivity. The within module connectivity of a gene is a measure of the strength of its co-expression with all other genes in the module to which it was assigned, whereas total connectivity is a gene's level of coexpression with all genes in the dataset. Genes that fell in the upper 0.95-quantile of within module connectivity (exceeding values of 298 and 139 kWithin for brain and liver, respectively) were treated as within module hub genes in downstream analysis, and genes that fell in the upper 0.95-quantile of total connectivity (exceeding values of 516 for brain and 354 for liver) were treated as overall high connectivity hub genes in downstream analysis. Based on these steps, we identified 600 and 607 intramodular (kWithin) hub genes and 705 and 639 hub genes with high overall connectivity (kTotal) for brain and liver, respectively.

To estimate the expected number of misexpressed hub genes by chance, we ran 1,000 simulations using the observed data. For each simulation, we randomly sampled the observed number of hub genes (for both kWithin and kTotal datasets) from the full set of genes in the RNAseq data and counted the number of randomly sampled genes that were misexpressed in $F_1$ hybrids. We then asked whether the observed number of hub genes that were misexpressed exceeded expectations from these null datasets.

Intriguingly, more hub genes were misexpressed in $F_1$ hybrids at ambient temperature in the brain than expected by chance (15 within module and 20 total connectivity hub genes; 95% CI of simulations = 2-12 genes), but none were misexpressed at high temperature (95% CI of simulations = 0-6 genes). These results suggest that more highly connected genes (i.e. those with more genetic interactions) are more likely to be constitutively misexpressed in $F_1$ hybrids than expected by chance. However, we did not observe an unexpected level of misexpression of hub genes in $F_1$s at high temperatures. We observed no misexpressed hub genes in the liver under either thermal context (as expected by chance based on simulation results; 95% CI of simulations = 0-3 for both within and total connectivity simulations).

*Supporting Information 7. Misexpression in other temperature-associated modules*

The misexpression patterns observed for circadian clock genes in $F_1$s (see main text) caused us to ask more generally about $F_1$ misexpression in the other 11 brain and one liver temperature-associated modules. We used permuted null distributions to build our expectations, as described in the previous section. We found that, overall, in the brain, two modules had fewer and two modules had more misexpressed genes at ambient temperature in $F_1$ hybrids than expected by chance (see Table S12). At high temperatures, one module had fewer and three modules had more misexpressed genes in $F_1$ hybrids than expected by chance (see Table S12). In the liver, only one module (greenyellow) had a higher than expected number of misexpressed genes at the high temperature treatment (though this module is not temperature-associated overall).

Of the four temperature-associated brain modules with more misexpressed genes than expected, each had ~5-10% of their gene members exhibiting misexpression in $F_1$s (compared to a maximum of 1-2% expected by chance; Table S12). The genes assigned to these modules were further evaluated with GO enrichment analysis to determine which biological pathways may be affected by misexpression in $F_1$s (see Methods).

One of the two brain modules enriched for $F_1$ misexpression at ambient temperature (grey60) is temperature-associated and highly enriched for immune system regulation and response genes, including signaling of *nfkb1*, the master regulator of the immune response. As discussed in the main text, these pathways were also enriched for differential expression in $F_1$ hybrids, but not *X. malinche* or *X. birchmanni*, based on KEGG analysis. The three modules with excess $F_1$ misexpression at high temperature were enriched for metabolic processes, which are commonly regulated in response to temperature, including in maintenance of homeostasis. Specifically, the module skyblue was enriched for pathways including primary, cellular, nucleic acid, and RNA metabolic processes. All misexpressed genes in the lightgreen module, including the circadian clock gene *cry1a,* fell into the GO categories 'cellular macromolecule metabolic process' and 'regulation of cellular process'.

*Supporting Information 8. Exploration of co-expression modules associated with both temperature and genotype*

WGCNA analysis identified several modules that were significantly associated with both temperature condition and genotype. Two temperature-associated brain modules (purple and red) were significantly associated with species (both *X. malinche* and *X. birchmanni*), one (skyblue) was associated with *X. malinche* alone, and one (paleturquoise) was associated with both *X. birchmanni* and $F_1$ genotypes (Table S11). GO analysis of these four brain modules indicated that several metabolic processes were enriched in the red, skyblue, and paleturquoise modules, while the genes in purple module were enriched for immune response and signaling pathways.

*Supporting Information 9. Additional discussion of expression patterns observed in the circadian clock pathway*

In the main text, we describe patterns of misregulation of genes in a circadian rhythm associated module in $F_1$ hybrids. Here, we highlight some particularly interesting patterns of expression of genes in this module. First, it is worth noting that in mammals, the molecular core of the circadian clock is composed of the CLOCK/ARNTL heterodimer transcription factor, three period (*per*) circadian regulator, and two cryptochrome (*cry*) circadian regulator genes. CLOCK/ARNTL activates transcription of *per* and *cry* genes, which in turn form complexes that inhibit CLOCK/ARNTL activity [23]. The cyclical oscillation of *per* and *cry* gene expression drives the circadian rhythm. Due to the whole genome duplication event in teleosts, most ray-finned fishes have two paralogs for each mammalian clock gene [24] and researchers have speculated that there may have been subsequent subfunctionalization or functional diversification in this pathway [25,26]. However, the core molecular clock pathway appears to be conserved between mammals and fish [27–31].

One gene identified in this module that is misexpressed in $F_1$ hybrids at both ambient and high temperature is the transcription factor *dbpb* (Fig. 3C). *dbpb* is a paralog of the mammalian *dbp* gene and appears to have aberrant expression in $F_1$ hybrids across all conditions studied here. In both the brain and liver, $F_1$s have low expression of *dbpb* compared to the expression levels observed in the parental species at ambient temperature conditions (22.5°C). Both *X. birchmanni* and *X. malinche* strongly downregulate expression of *dbpb* in response to high temperature, whereas $F_1$s fail to change *dbpb* expression, leading to high expression of *dbpb* in $F_1$s relative to the parental species at 33.5°C in both tissues (Fig. 3C). Misexpression of *dbpb* in both thermal contexts and across tissues suggests that this gene may be broadly misregulated in *X. malinche-X. birchmanni* $F_1$ hybrids.

Because *dbpb* is a transcriptional driver of one of two CLOCK/ARNTL-mediated circadian clock feedback loops, we were interested in whether the genes it is known to regulate were also misexpressed in $F_1$ hybrids. All period genes (*per1*, *per2*, *per3*), nuclear receptor 1d (*nr1d1/nr1d2*, the orthologs of *rev-erb alpha/beta*) genes, and RAR-related orphan receptor (*ror*) genes have D-box elements in their promoters which are activated by *dbpb* [32]. We looked for patterns of misregulation in these *dbpb*-controlled clock genes. While expression of the *ror* genes does not seem to be impacted by *dbpb* misexpression, all three *per* genes and *nr1d2a/b* are also strongly misexpressed in $F_1$s, suggesting that misexpression of *dbpb* results in a domino effect in misexpression of downstream target genes. The consequences of these kinds of cascading networks of misexpressed genes is an exciting avenue for future work understanding the fitness of hybrids.

Intriguingly, *nr1d1* is involved in a regulatory feedback loop with *dbpb* and degrades under stress to relax inhibition of the inflammatory response [33]. Notably, *nr1d1* is more highly expressed in *X. malinche* across temperature conditions. This might suggest that the inflammatory response mediated by *nr1d1* is constitutively more active in *X. birchmanni* than in *X. malinche*, consistent with the observation that some negative regulators of the inflammatory response, such as *tnfaip3* and *nfkbia*, are strongly downregulated in *X. birchmanni* in response to high temperature (Fig. S8; as discussed in Supporting Information 5).

Finally, though mRNA levels provide some insight into clock regulation, we note that gene expression data provides only a partial picture of the regulatory landscape of circadian processes, which are well-known to be modulated by posttranslational regulation [34,35]. Though the expression data presented here clearly supports hybrid dysfunction at the expression-level, there may be further breakdown, or alternatively, compensations, of clock protein levels [36].

*Supporting Information 10. Simulations evaluating clock gene ancestry in natural hybrid zones*

The interacting clock genes *nr1d2b* and *arntl1a*, both of which are misexpressed in $F_1$ hybrids, share exceptionally high *X. birchmanni* ancestry in both the Acuapa (99% and 92%, respectively) and Tlatemaco (90% and 89%, respectively) natural hybrid populations. To evaluate the probability of a pair of genes sharing such high *X. birchmanni* ancestry across these two populations, we calculated the average *X. birchmanni* ancestry in the two populations for every gene across the genome, and randomly selected 10,000 pairs of genes. For each pair, we determined whether both genes had *X. birchmanni* ancestry above that observed for the focal genes in the real data (e.g. 99% for Acuapa and 90% for Tlatemaco for *nr1d2b* ancestry). Of all 10,000 randomly sampled pairs of genes, the probability of both genes having ancestry proportions above that threshold was 1.9% in Acuapa and <0.01% in Tlatemaco. We also subsetted pairs where at least one of the two genes exceeded the ancestry threshold and estimated the probability of observing this level of *X. birchmanni* ancestry based on this criterion to be 7.0% for Acuapa and <0.01% for Tlatemaco. These simulations indicate that the level of *X. birchmanni* ancestry in *nr1d2b* and *arntl1a* in both populations is unlikely to be due to chance, hinting that there may have been selection favoring *X. birchmanni* ancestry at these loci in natural populations.

*Supporting Information 11. Relationship between $CT_{max}$ and the melanoma hybrid incompatibility*

A clear pattern that emerged from our analyses was that hybrids often operated outside of parental ranges when it came to particular traits (e.g. gene expression patterns, response to thermal stress based on chromosome 22 genotype). Previous work in our group has documented the presence of hybrid incompatibilities between *X. malinche-X. birchmanni* that impact physiological traits. In particular, we were interested in exploring how a hybrid incompatibility that generates melanoma interplays with thermotolerance (Fig. 4B).

While we cannot directly evaluate this claim with existing data, we speculate that the observed relationship between the presence of melanoma and lowered $CT_{max}$ could be driven by compounded stress and/or taxed stress response machinery. Presumably, fish with cancer experience elevated stress levels in their baseline physiological function and existing data hints that they also have compromised immune responses [37,38]. While hybrid individuals with even early stage melanoma have reduced $CT_{max}$, hybrid individuals with a normal spotting phenotype perform just as well as individuals that lack the spot. This indicates that the reduction in $CT_{max}$ is related to the presence of melanoma, rather than pleiotropy in genes underlying the spotting pattern itself (as is the case for micromelanophore pigmentation patterns and $CT_{max}$ in *X. variatus*, [39]). Exploring the range of physiological impacts of the melanoma hybrid incompatibility and its links to thermotolerance in natural hybrid populations is an exciting area for future work.

*Supporting Information 12. $CT_{max}$ as a measure of thermotolerance*

In the main text, we treat our experimental measure of thermotolerance, the critical thermal maximum or $CT_{max}$, interchangeably with thermotolerance. However, "thermal tolerance" encompasses a wide range of traits. It is likely that the genetic architecture underlying these traits overlap but are not completely shared. The $CT_{max}$ trait is widely used to measure the upper thermal tolerance limit in ectotherms and is therefore useful in that it is generally comparable between the present study and past studies. Specifically, this metric is ecologically-meaningful, and a good predictor of the temperatures individuals from natural populations can withstand in the short-term without experiencing mortality [40]. However, it is important to note that $CT_{max}$ measures the tolerance of an ectotherm to acute heat stress, as opposed to long-term shifts in thermal environment. Therefore, it is possible that an individual's $CT_{max}$ may have modest predictive power of its tolerance to an increase in temperature over longer time periods. Investigating the links between $CT_{max}$ and long-term thermal resilience in swordtails may be worth exploring for mapping studies in the future.

*Appendix 1. Key commands used in analysis*

*kallisto*
To obtain transcript counts from trimmed RNAseq read data, we built an index for the *X. birchmanni* reference transcriptome with `kallisto index` and then pseudoaligned reads to this reference with `kallisto quant`.

```
kallisto index -i xbirch_transcriptome.fa
kallisto quant -i xbirch_transcriptome.idx -o S1_TSR1-22C-Xmal-brain_1-A1_kallisto
S1_TSR1-22C-Xmal-brain_1-A1_R1_val_1.fq.gz S1_TSR1-22C-Xmal-brain_1-
A1_R2_val_2.fq.gz
```

*R/qtl*
To map $CT_{max}$ QTL, markers across the genome were thinned and filtered (see Methods), and recombination rate was estimated between markers with `est.rf` and genotype probabilities were calculated with `calc.genoprob`. Covariates were selected as described in the Methods and stored in a covariates vector. A single-scan for QTL was performed with scanone with a model including the covariates, added using the 'addcovar' argument. Finally, to calculate the significant genome-wide LOD threshold, we used 1,000 permutations, where single QTL scans were performed on marker data matched to randomly shuffled $CT_{max}$ and covariate phenotypes 1,000 times. See Methods for more information and find the full pipeline on GitHub at https://github.com/Schumerlab/thermotolerance.

```
data_sub.ds = est.rf(data_sub.ds)
data_prob = calc.genoprob(data_sub.ds)
covariates = pull.pheno(data_prob, c("hybrid_index","STH.1","STH.2","STH.3","STH.4",
                                     "STH.7","STH.8","STL.1","STL.2","STL.4","STL.5",
                                     "STL.6","STL.7","STM.2", "STM.4","STM.5","STM.6",
                                     "STM.7"))
scanone(data_prob, method="hk",addcovar=covariates)
scanone(data_prob, method="hk", addcovar=covariates, n.perm=1000)
```

*WGCNA*
To construct a weighted, single-block, unsigned network, we used a topological overlap matrix, a minimum module size of 20, maximum block size of 20,000 (to capture all genes in the genome in one block), block size penalty power of Infinity (since maxBlockSize > total genes), no reassignment (reassignThreshold = 0), module merge cut height of 0.25, pamStage/pamRespectsDendro FALSE, and default values for all other parameters. See full script on GitHub at https://github.com/Schumerlab/thermotolerance.

```
blockwiseModules(
         datExpr, power = 7,
         TOMType = "unsigned", minModuleSize = 20,
        maxBlockSize = 20000, blockSizePenaltyPower = Inf,
        reassignThreshold = 0, mergeCutHeight = 0.25,
        numericLabels = FALSE, pamRespectsDendro = FALSE,
```

```
loadTOMs = FALSE,
saveTOMFileBase = "TT-Brain-TOM",
verbose = 3)
```

**Supplementary Figures**

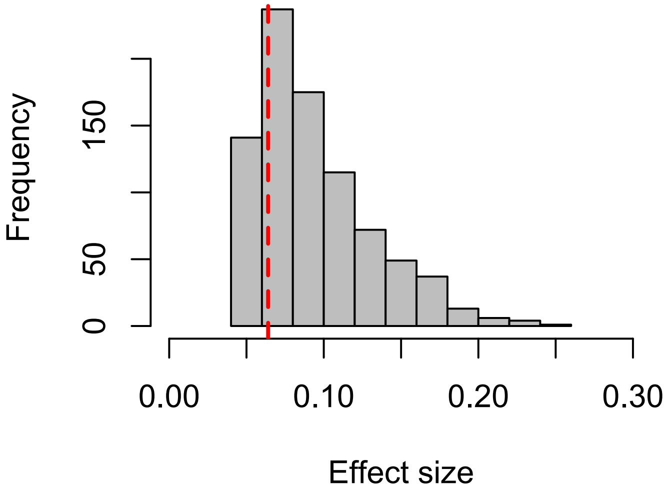


**Figure S1**. Posterior distribution of effect sizes for the chromosome 22 QTL based on 500 accepted simulations from ABC (see Supporting Information 1 for more information). The maximum *a posteriori* estimate from the accepted simulations is represented by the red dotted line.

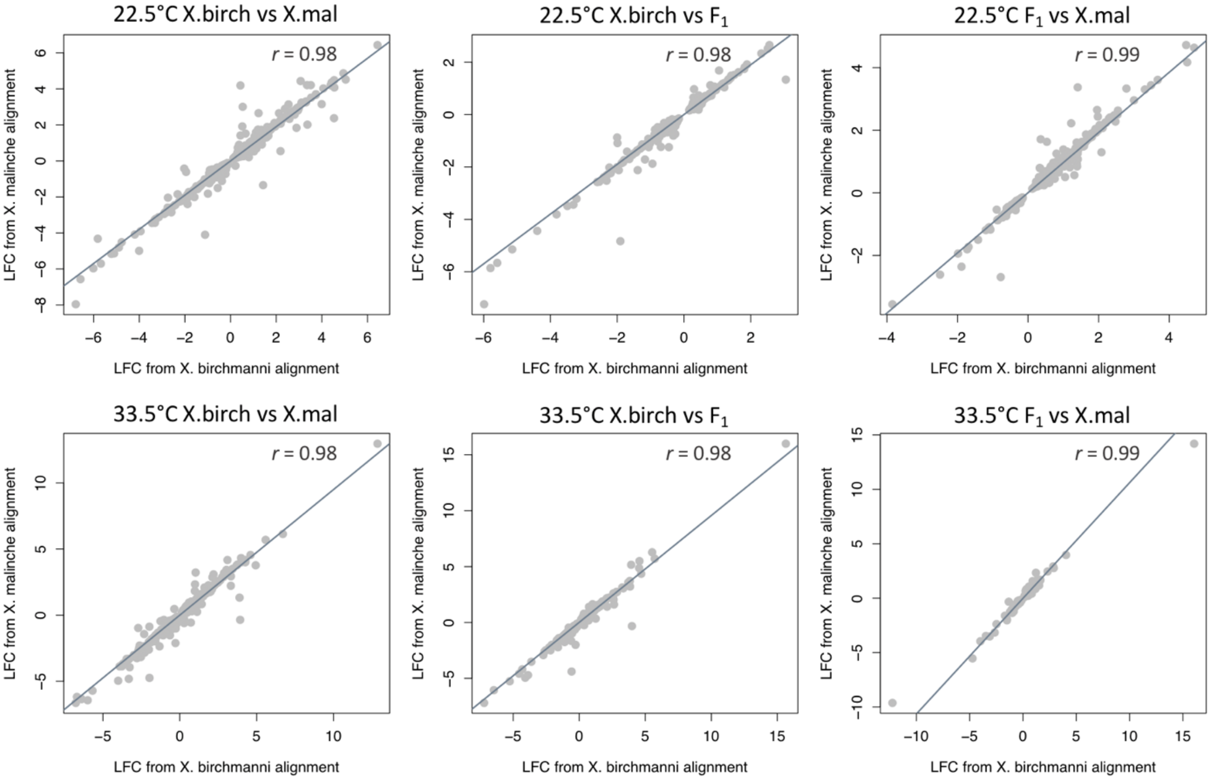


**Figure S2**. Plots showing correlations of log-fold change (LFC) values based on analyses using the *X. birchmanni* versus *X. malinche* reference transcriptomes for pseudoalignment of brain RNAseq reads for different comparisons. Each grey point represents one gene's LFC values calculated from each reference. Pearson's correlation coefficients (*r*) for each comparison are provided in the inset text.

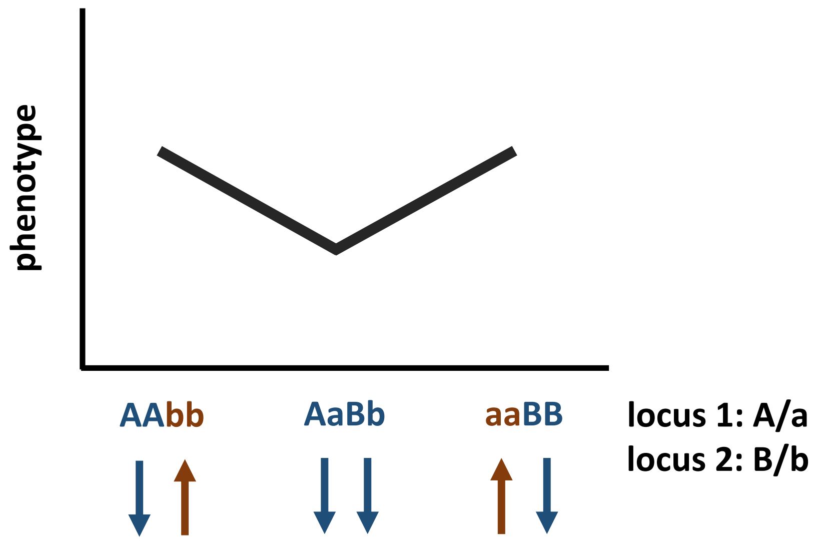


**Figure S3**. Schematic of pseudo-underdominance. Two loci with opposing effects on phenotype are linked, and one allele of each (*A* and *B*) is dominant over the other (*a* and *b*, respectively). The effect of the genotypes on phenotype is denoted by blue and red colored arrows. In this case, the effects of the linked loci neutralize each other in homozygotes. However, in heterozygotes, dominance effects at both loci result in a compounded reduction in phenotype, resulting in a phenotype profile resembling that expected from underdominance at a single locus.

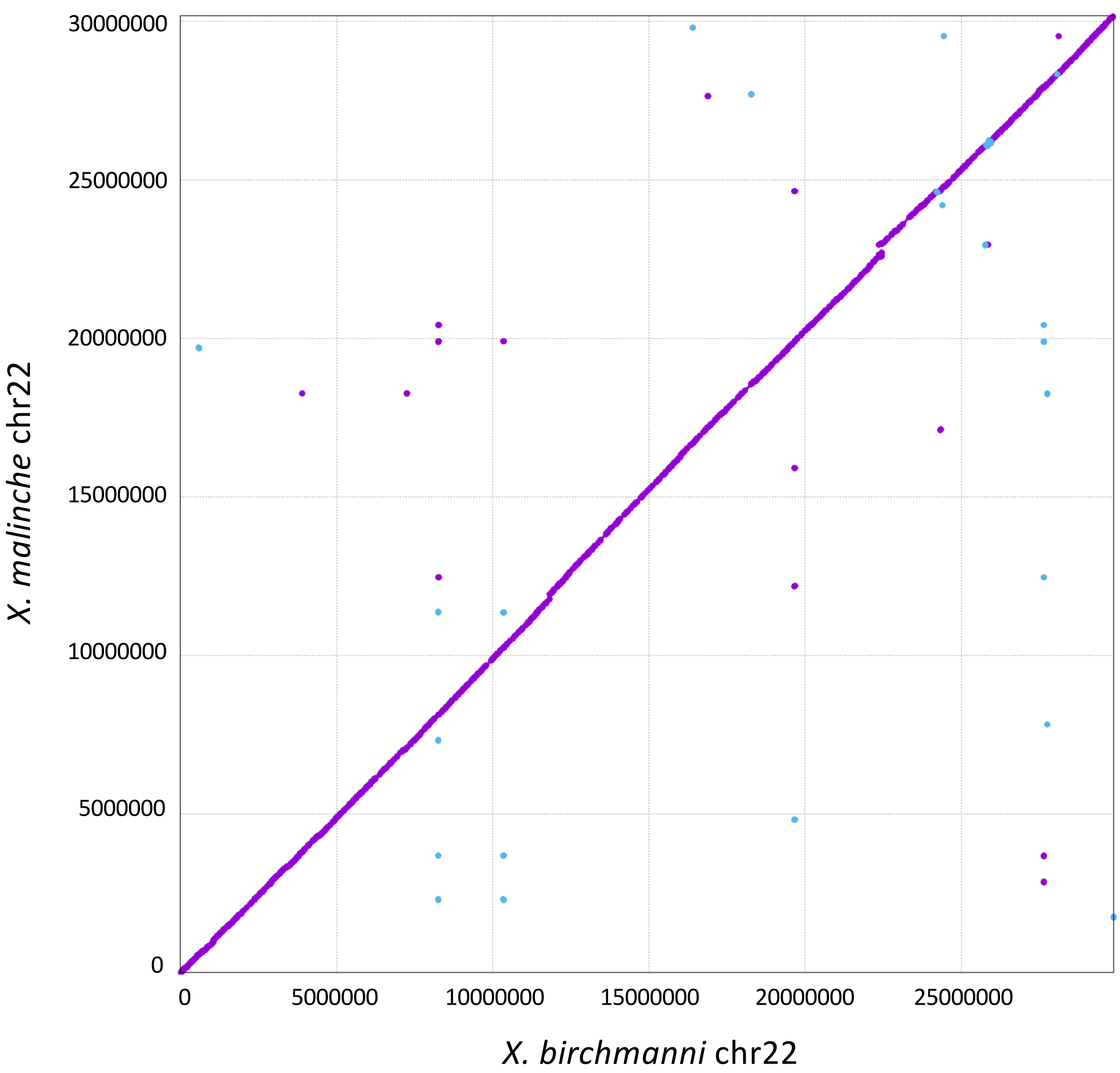


**Figure S4**. Alignment of the *X. birchmanni* and *X. malinche* chromosome 22 PacBio assemblies to rule out differences in chromosomal structure (such as inversions) as an explanation for the signal observed at the chromosome 22 $CT_{max}$ QTL. Alignments were produced using MUMmer4 [11].

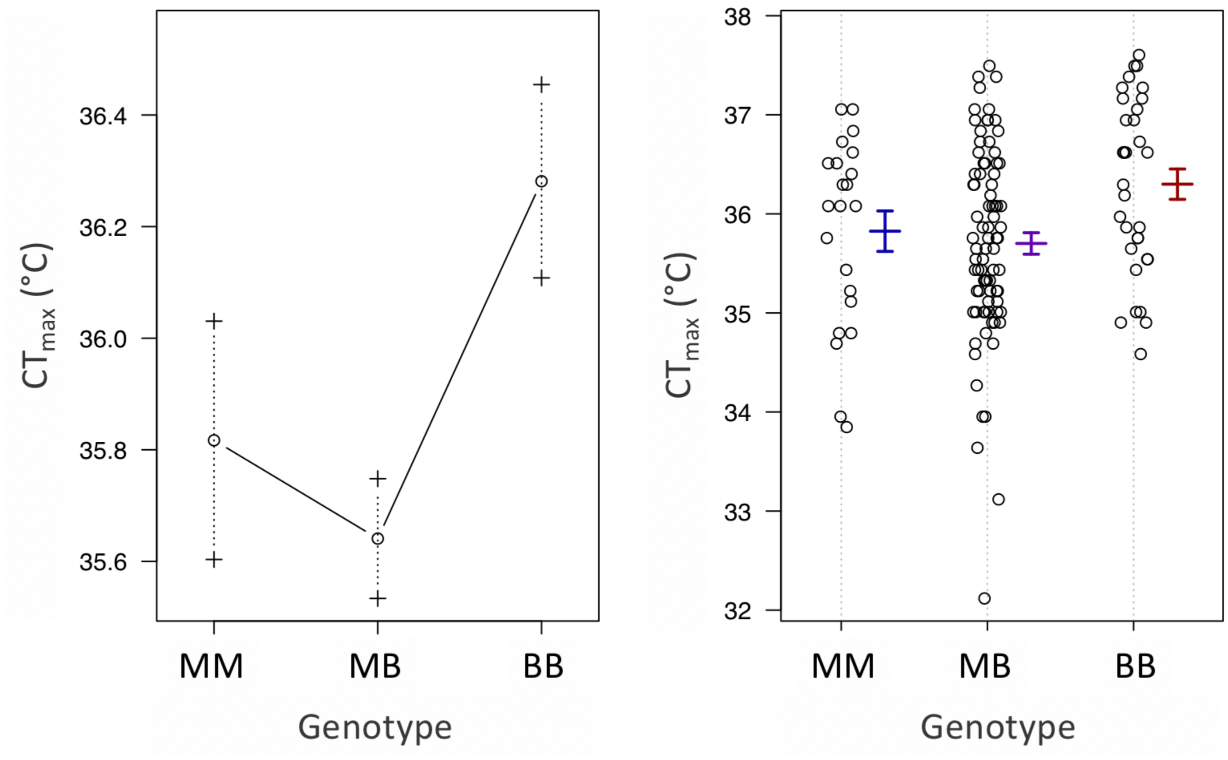


**Figure S5**. Two plots showing the effect of individual genotypes at the chromosome 15 QTL peak (~4.4Mb) on $CT_{max}$ phenotype. Plotted on the left are the average $CT_{max}$ values for each genotype and error bars showing one standard error for each group. On the right, the same means and error bars are plotted but the $CT_{max}$ data of individual artificial hybrids is also shown.

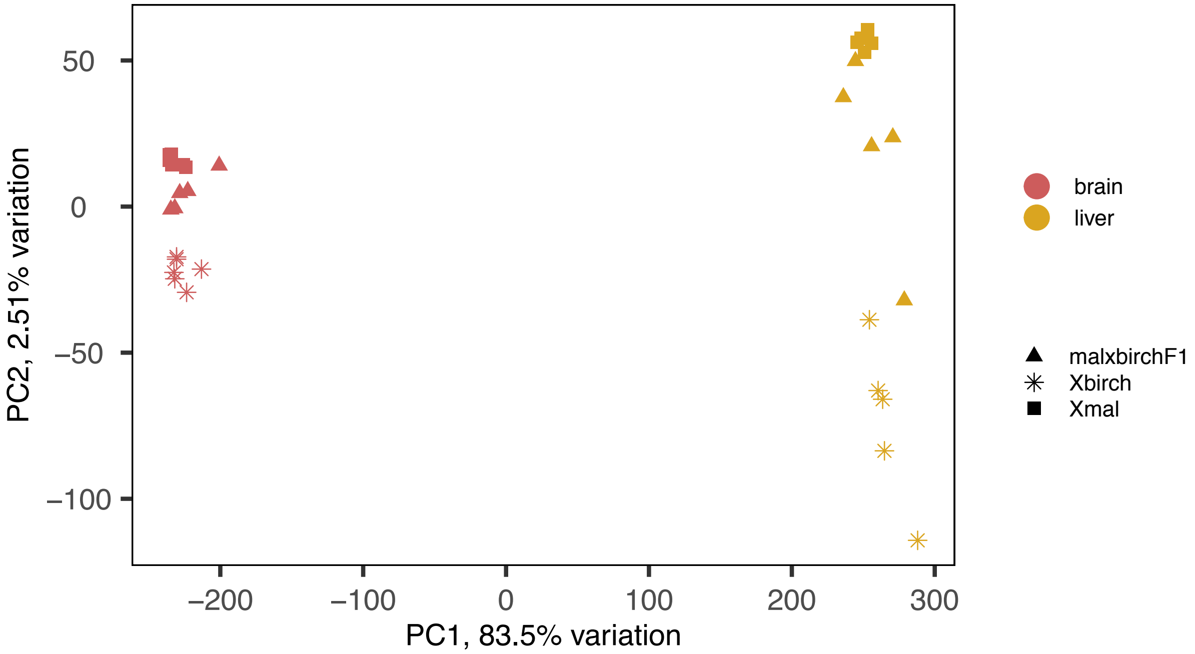


**Figure S6.** PCA of RNAseq data including both brain and liver datasets. In this analysis, tissue type explains the vast majority of variation in expression observed across these datasets (PC1, 83.5%).

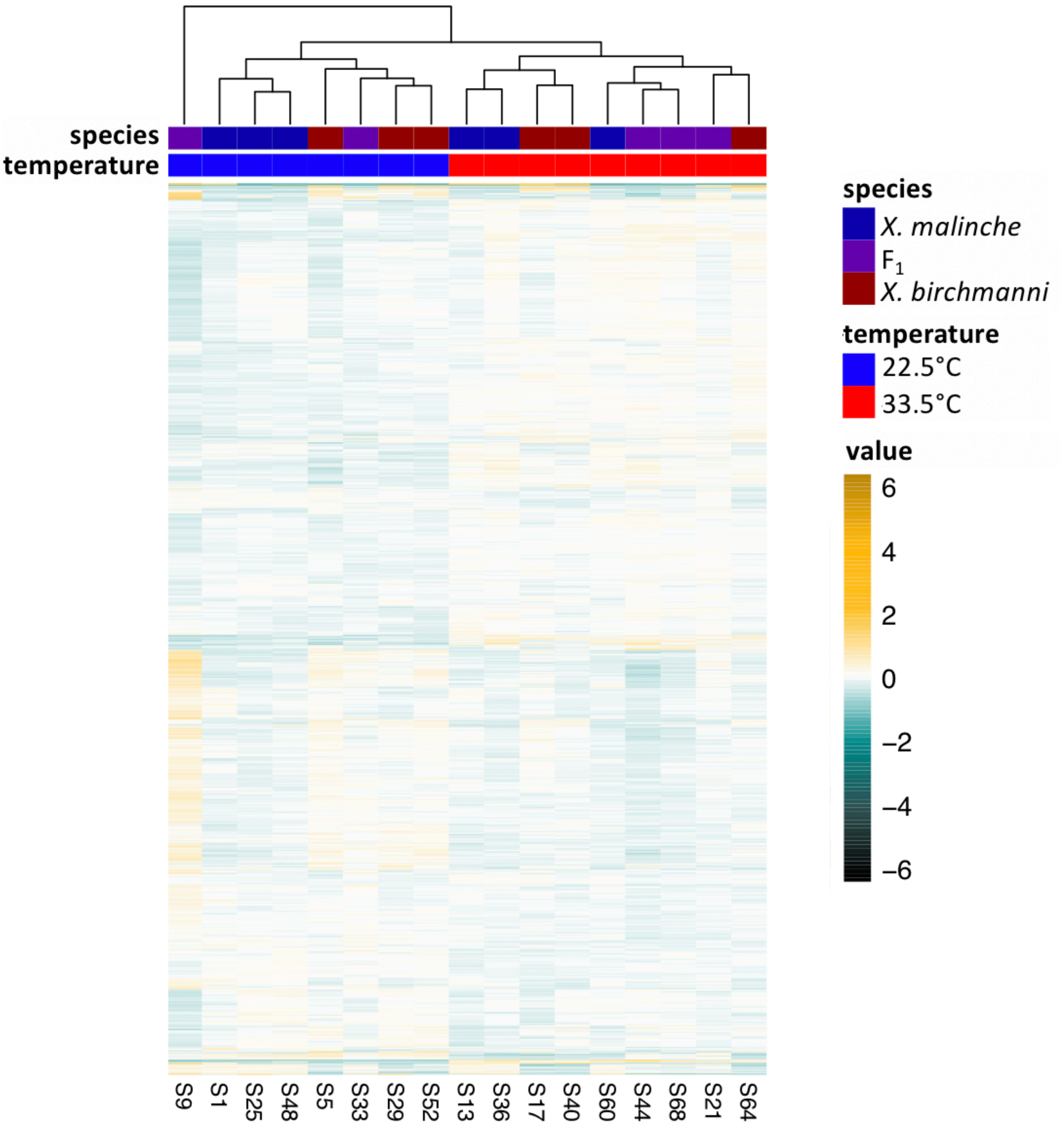


**Figure S7**. Heatmap of all genes that are significantly differentially expressed between temperature treatments for at least one comparison in *X. malinche*, *X. birchmanni,* and $F_1$ brain tissue (3,349 genes total). Samples are clustered on the x-axis and genes are clustered on the y-axis by Euclidean distance. Each row represents one gene, and each cell represents the raw distance of the corresponding sample's normalized gene count from the mean of counts across samples for that gene. Normalized gene counts were transformed to have approximately constant variance across samples (using the DESeq2 function `varianceStabilizingTransformation`). Value colors represent the transformed counts minus the mean of that gene's counts across samples.

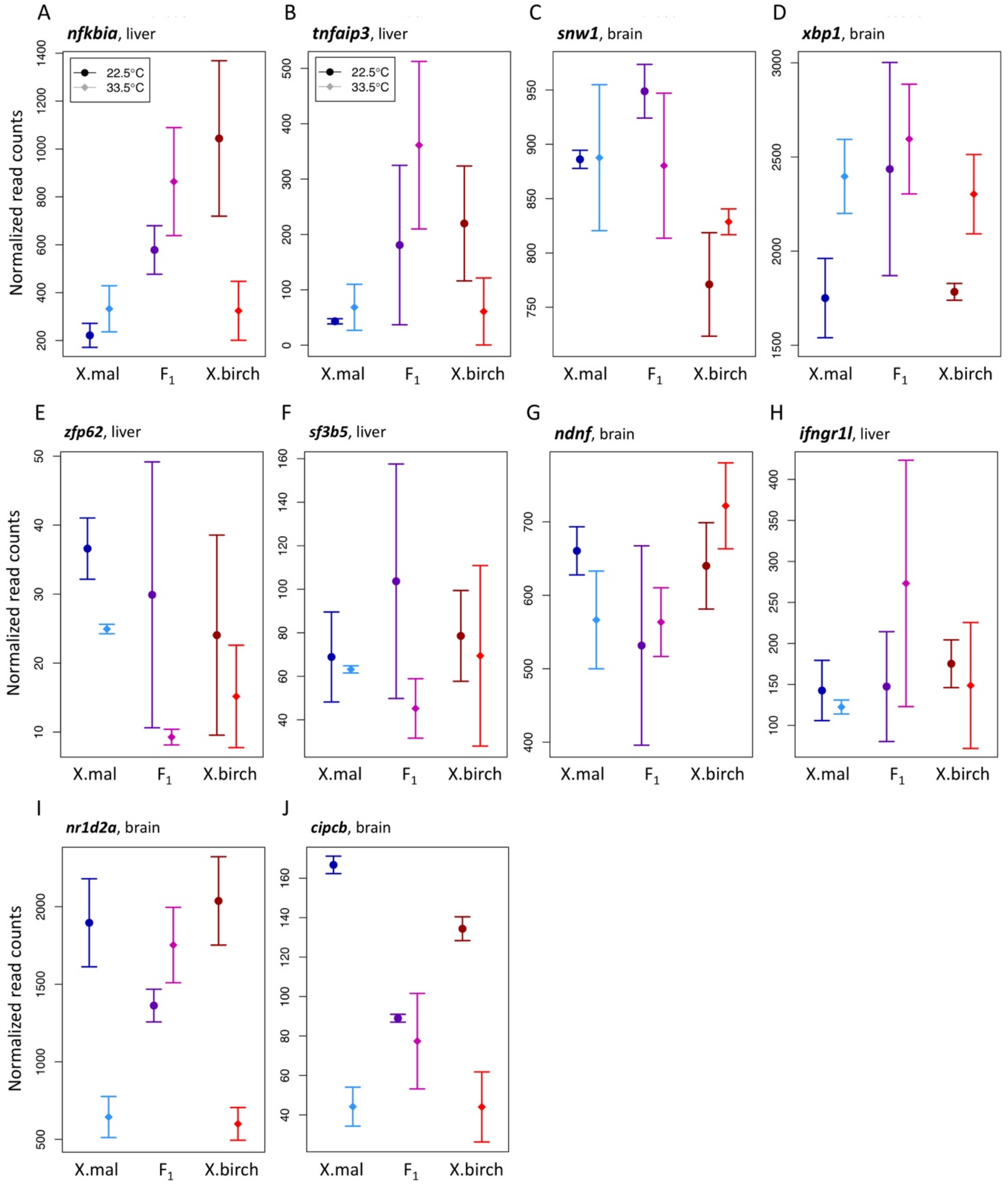


**Figure S8**. Expression plots for **A**) *nfkbia* in liver, **B**) *tnfaip3* in liver, **C***) snw1* in brain, **D**) *xbp1* in brain, **E**) *zfp62* in liver, **F**) *sf3b5* in liver, **G**) *ndnf* in brain, **H**) *ifngr1l* in liver, **I**) *nr1d2a* in brain, and **J**) *cipcb* in brain. Mean normalized counts at 22.5°C are represented by a circle in a darker color and mean normalized counts at 33.5°C are represented by a diamond in a brighter color. Error bars show one standard deviation of expression.

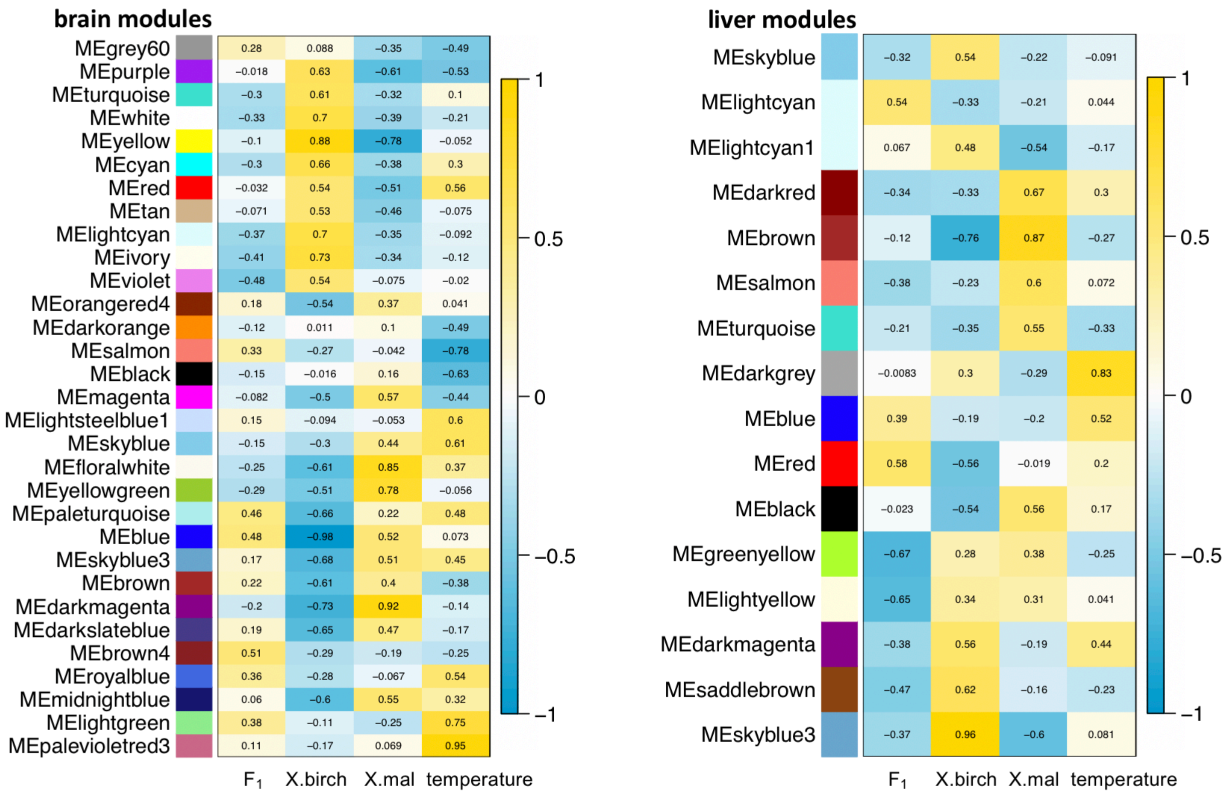


**Figure S9**. Weighted gene co-expression analysis uncovered 12 temperature-associated modules in the brain (Fig. 3B) and 2 in the liver, in addition to several other modules associated with genotype. Shown here are correlation heatmaps for the modules significantly associated (p-value < 0.05) with at least one trait of interest ($F_1$, *X. birchmanni*, or *X. malinche* genotype or temperature) in the brain and liver RNAseq datasets. Traits are listed on the x-axis, and color blocks and labels on the y-axis represent the WGCNA module. Pearson's correlation coefficients are listed for each module and trait, with box color corresponding to the strength of the correlation (yellow spectrum for a positive trait-module correlation, blue spectrum for a negative trait-module correlation). Note that because module names are arbitrarily assigned in each analysis, shared module names between the brain and liver datasets do not represent the same gene members or gene cluster.

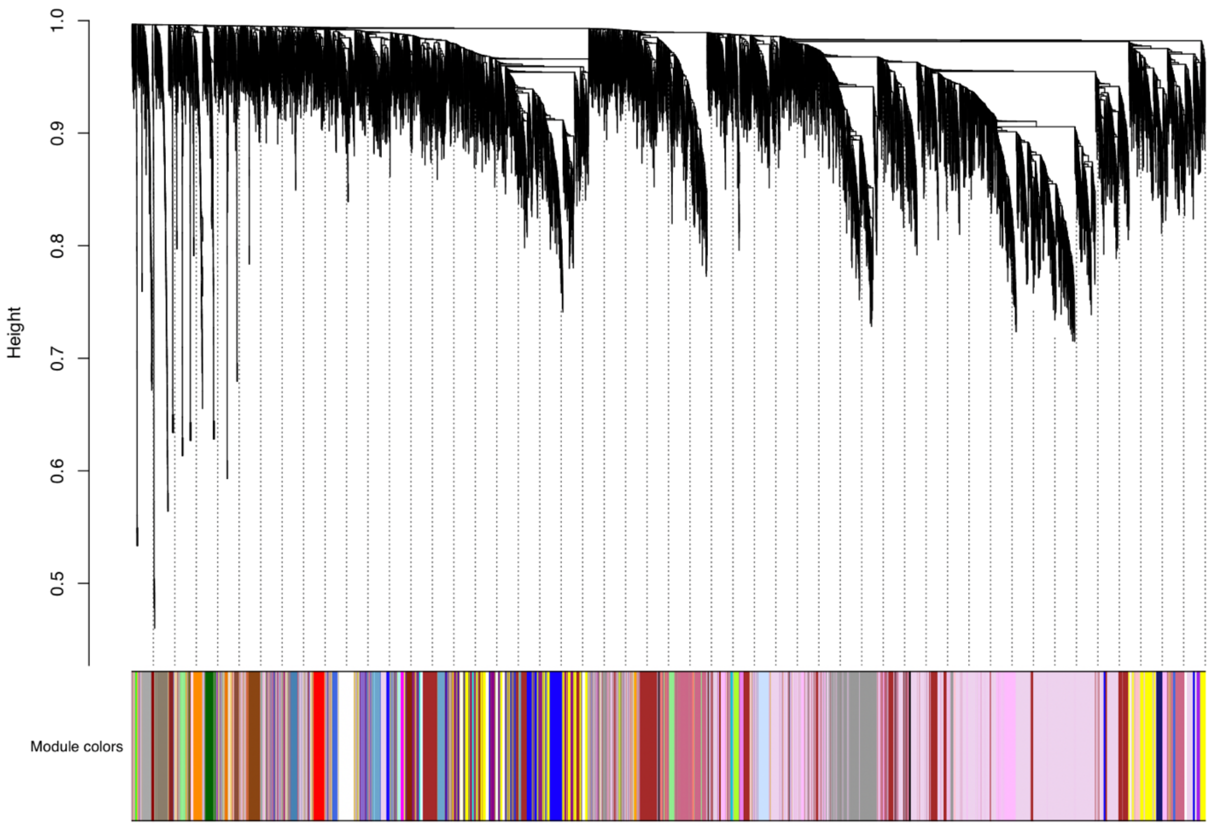


**Figure S10**. Dendrogram of WGCNA hierarchical gene clustering of brain RNAseq data. Branches represent individual genes, and the color panel indicates the modules that genes were grouped into, based on WGCNA's co-expression clustering algorithm.

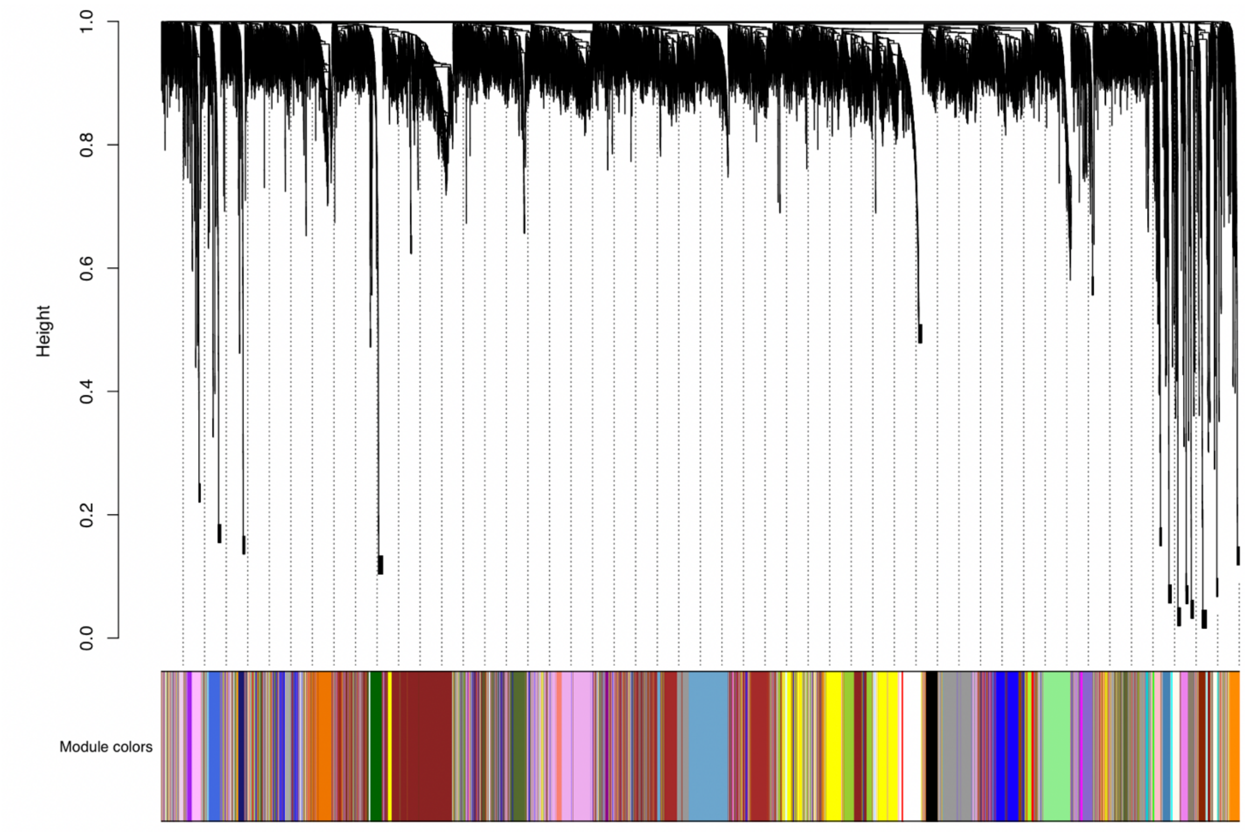


**Figure S11**. Dendrogram of WGCNA hierarchical gene clustering of liver RNAseq data. Branches represent individual genes, and the color panel indicates the modules that genes were grouped into, based on WGCNA's co-expression clustering algorithms.
